# Viral route of infection determines the effect of *Drosophila melanogaster* gut bacteria on host resistance and tolerance to disease

**DOI:** 10.1101/2021.02.18.431843

**Authors:** Miguel Landum, Marta Salvado Silva, Nelson Martins, Luís Teixeira

## Abstract

The microbial community interacting with a host can modulate the outcome of pathogenic infections. For instance, *Wolbachia*, one of the most prevalent invertebrate endosymbionts, strongly increases resistance of *Drosophila melanogaster* and other insect hosts, to many RNA viruses. *D. melanogaster* is also in continuous association with gut bacteria, whose role in antiviral immunity is poorly characterized. Here we asked how gut-colonizing bacteria impact viral titres and host survival, and how these interact with route of infection or *Wolbachia* presence. We compared germ-free flies and flies associated with two gut bacteria species recently isolated from wild flies (*Acetobacter thailandicus and Lactobacillus brevis*). We found that *Wolbachia*-conferred protection to both DCV or FHV is not affected by the presence or absence of these gut bacteria. Flies carrying *A. thailandicus* have lower DCV loads than germ-free flies, upon systemic infection, but reduced survival, indicating that these bacteria increase resistance to virus and decrease disease tolerance. Association with *L. brevis*, alone or in combination with *A. thailandicus*, did not lead to changes in survival to systemic infection. In contrast to the effect on systemic infection, we did not observe an impact of these bacteria on survival or viral loads after oral infection. Overall, the impact of gut-associated bacteria in resistance and tolerance to viruses was mild, when compared with *Wolbachia*. These results indicate that the effect of gut-associated bacteria to different viral infections, and different routes of infection, is complex and understanding it requires a detailed characterization of several parameters of infection.

## Introduction

Insects, as many animal and plants, host a wide variety of microbes. These interactions influence differently host biology, with a wide range of deleterious and beneficial impacts on several host traits (Dillon and Dillon 2004; Engel and Moran 2013). One common impact of this microbial community is the modulation of host-pathogen interactions.

A particularly well-studied case is the association of *Wolbachia* and its hosts. *Wolbachia* is the most prevalent intracellular symbiont of insects being present in 40% of insect species and is known to manipulate several host traits (Zug and Hammerstein 2012; Werren, Baldo, and Clark 2008). One of the outcomes of this interaction is the ability of *Wolbachia* to confer host resistance against several RNA viruses that infect *Drosophila* and mosquitoes (Teixeira, Ferreira, and Ashburner 2008; Hedges et al. 2008; Moreira et al. 2009).

Gut-associated microbes have also been shown to play an essential role in restricting pathogen proliferation and their detrimental outcomes in insects (Dong, Manfredini, and Dimopoulos 2009; Koch and Schmid-Hempel 2011). The protective role of gut microbes has been established against some groups of pathogens, and its interactions with viruses has been explored in some systems. In the mosquito *Aedes aegypti*, antibiotic treatment leads to depletion of gut microbiota, decrease in basal Toll pathway activity and increase in dengue viral loads in the gut upon infection (Xi, Ramirez, and Dimopoulos 2008). Furthermore, the mono-association of *A. aegypti* with certain bacteria isolated from wild-caught mosquitos had increased expression of antimicrobial peptides (AMPs) and lower titres of dengue virus (Ramirez et al. 2012). These results indicate that, in mosquitoes, the gut microbiota is modulating resistance to viruses, through the basal activation of gut immunity. However, a different study report that an infection of the same mosquito specie with Serratia, increased its susceptibility to dengue virus (Apte-Deshpande et al. 2012). These contrasting results suggest an intricate and species specific effect on immunity and virus inhibition of gut bacteria.

*Drosophila melanogaster* is also in continuous association with bacteria in the gut, either acquired through feeding or colonizing it (Pais et al. 2018; Broderick and Lemaitre 2012). Presence of some members of the gut-associated bacteria reduce the viral titres in the gut of orally infected flies (Sansone et al. 2015). As in mosquitoes this is mediated by priming of basal immunity in the gut (Sansone et al. 2015). On the other hand, gut-associated bacteria do not seem to affect systemic viral infection outcome or interact with *Wolbachia* in this setup (Ye et al. 2017). Therefore, the interaction of gut bacteria with viral infection is also complex and seems to depend on bacterial species and route of infection.

Here, we tested the modulation of viral infections by gut-associated bacteria recently isolated from wild flies (Pais et al. 2018). These isolates of *Acetobacter thailandicus* and *Lactobacillus brevis* can proliferate and colonize the gut of D. melanogaster and, therefore, interact with the host differently than most lab associated bacteria strains (Pais et al. 2018). We tested their influence on infection with different viruses, different routes of infection, and in the presence and absence of *Wolbachia*. Moreover we determined the outcome of viral infection by analysing survival and viral titres in order to better assess their impact on resistance and disease tolerance (Soares, Teixeira, and Moita 2017). This set of data allows a broad comparative view on how gut colonizing bacteria influence viral infection outcome in *D. melanogaster*.

## Materials and Methods

### Fly strains and husbandry

*Drosophila melanogaster w*^*1118*^ isogenic background (Ryder et al. 2004) carrying *Wolbachia w*MelCS_b (Wolb^+^) or not (Wolb^-^) was used in most of the experiments. For systemic infections of mono-associated flies, a different host genetic background (Oregon R W20) with and without a wMel-like *Wolbachia*, was also used (Teixeira, Ferreira, and Ashburner 2008; Chrostek et al. 2020).

Flies were raised and experiments performed on typical autoclaved VDRC Vienna food with minor adaptations: 8g agar, 80g molasses, 22g beet syrup, 10g soy flour, 80g cornmeal, 18g yeast, 1.1L of water and 2.6% antifungal mix (0.2g Methyl 2-benzimidazolecarbamate, 100g methylparaben in 1L absolute ethanol).

### Axenic and Gnotobiotic (mono/di-associated) flies

Germ-free flies were raised as described by Pais et *al*. (Pais et al. 2018). Briefly, four to six hours-old embryos were collected and dechorionated using a 2.1% sodium hypochlorite solution, followed by washing with 70% ethanol and rinsed with sterile water. Embryos were then transferred to germ-free Vienna food, under sterile conditions.

To raise gnotobiotic flies, germ-free embryos were placed in vials containing Vienna food and 40µL of an overnight culture of *Acetobacter thailandicus* (isolate 1153/12) or *Lactobacillus brevis* (isolate 0356/12) were added, for mono-associated flies, and 20µL of A. *thailandicus* plus 20µL of L. brevis in the case of di-associated flies. Gnotobiotic flies were always raised one day after germ-free flies due to a developmental delay that occurs when flies are raised in the absence of gut microbiota (Pais et al. 2018).

Axenic and gnotobiotic flies were maintained at a constant temperature of 25^°^ C, under a 16h:8h light:dark cycle.

To check the status of the flies (germ-free or associated with the specific microbiota), adults were plated in MRS (DeMan-Rogosa-Sharpe) agar medium on the day of collection and bacterial species were identified by colony morphology (Pais et al. 2018).

### Infections and Virus Production

All systemic and oral infections were performed in a sterile vertical flow hood, and all the material used was sterilized by autoclave.

Viruses were produced and titrated as in Ferreira et al. (Ferreira et al. 2014). Viral suspension was filtered using Acrodisc® Syringe 0.2µm Filters.

For infection, flies were anesthetized using CO_2_ and pricked in the dorsolateral thorax using 0.15 mm diameter needles (Austerlitz Insect Pins; FST# 26002-15) dipped in the virus solution. Infections were performed in 3-6 days old adult males or females. After viral or mock infection (25 flies per condition), 5 flies per vial were maintained in germ-free Vienna food and survival checked daily. Drosophila C Virus (DCV; 10^11^ TCID_50_/mL) infected flies were maintained at 18°C, while Flock House virus (FHV; 10^8^ TCID_50_/mL) infected flies were maintained at 25°C.

For oral DCV infection, ten 3-6 days old male or female flies, were put in contact with virus-soaked filter paper (200µL of a 10^11^ TCID_50_/mL viral suspension with 5% sucrose in each vial) for 24 hours. The flies were then transferred to germ-free Vienna food, kept at 25°C and survival checked daily. Systemic and oral infection experiments were repeated independently three times.

### RNA Extractions, cDNA Synthesis and Real-Time Quantitative PCR

RNA was extracted from single flies using TripleXtractor reagent (Grisp) according to the manufacturers’ instructions. Each sample was treated with DNase I (Promega) to eliminate possible DNA contaminations. RNA concentrations and purity were determined using NanoDrop ND-1000 Spectrophotometer.

cDNA was prepared from 0.1 µg of total RNA using Random Primers (Promega) and M-MLV Reverse Transcriptase (Promega). Primers were allowed to bind to the template RNA for 5 min at 70oC, followed by 25°C for 10 min, 37°C for 60 min and 85°C for 10 min.

For each RT-PCR reaction, 6 μL of iTaq SYBR Green supermix (Bio Rad), 0.5 μL of each primer solution at 3.6 mM and 5 μL of diluted DNA was used in a 384-well plate. Each plate contained two technical replicates of every sample for each set of primers. Primers used and viral RNA quantification using the Pfaffl method was described in Ferreira et al. (2014; Pfaffl 2001). RpL32 was used as a reference gene. Primers used were: Rpl32 forward 5’-CCGCTTCAAGGGACAGTATC-3’; Rpl32 reverse 5’-CAATCTCCTTGCGCTTCTTG-3’; DCV forward 5’-TCATCGGTATGCACATTGCT-3’; DCV reverse 5’-CGCATAACCATGCTCTTCTG-3’; FHV forward 5’-ACCTCGATGGCAGGGTTT-3’; FHV reverse 5’-CTTGAACCATGGCCTTTTG-3’. Real-time qPCR reactions were performed using the QuantStudio™ 7 Flex Real-Time PCR System (Applied Biosystems).

## Statistical analysis

Data analysis was performed using R software version 4.0.3 (Team 2012).

Survival analysis was done using a mixed effects Cox proportional hazard model, with sex and presence of each bacterial species (*Wolbachia, A. thailandicus or L. brevis*) as fixed factors and experimental replicate and individual fly vial as random variables, using the package coxme version 2.2-16 (Therneau 2020).

Differences on viral levels were tested using linear mixed effect models analysis on the log10 transformed viral RNA levels and with presence of each bacterial species (*Wolbachia, A. thailandicus or L. brevis*) as fixed factors and experimental replicate as random variable, using the lme4 version 1.1-26 package (Bates et al. 2015), or with a non-parametric Kruskal-Walis test when residuals from the linear modelling approach deviated strongly from normality, using the function kruskal.test in base R.

Multiple comparison analysis for both survival and viral titres were performed using the package emmeans version 1.5.3. (Lenth 2020). Survival curves were calculated using survminer package version 0.4.8. (Kassambara, Kosinski, and Biecek 2020). Graphics were generated using the ggplot package version 3.3.3, part of the tidyverse package version, 1.3.0. (Wickham et al. 2019).

The data and script for the analysis are available as supplementary data.

## Results

### Differences in disease tolerance and resistance to systemic viral infection in controlled *Drosophila*-bacteria interaction scenarios

We first tested the effect of the gut colonizing bacteria A. *thailandicus* on systemic infection with *Drosophila* C virus by comparing axenic and mono associated flies. We performed this experiment in lines with and without *Wolbachia*, to determine if *A. thailandicus* modulated this endosymbiont anti-viral protection, assessing survival to infection and viral loads (Fig 1 A-C). We saw no effect on presence or absence of this gut bacteria on *Wolbachia* protection, either in terms of survival (Mixed effects Cox model [COXME], *Wolbachia* X A. thailandicus effect, p = 0.93; Fig 1A, B) or viral titres (Linear mixed model [LMM], A. *thailandicus* X *Wolbachia, p* = 0.093, Fig 1C). Wolbachia presence increased survival upon viral infection (COXME, *Wolbachia* effect, *p* < 0.001, Fig 1A, B) and decreased viral titres by 1862-fold (LMM, Wolbachia effect, p < 0.001, Fig 1C), as expected (Teixeira, Ferreira, and Ashburner 2008; Hedges et al. 2008). On the other hand, the presence of the gut bacterium *A. thailandicus* had a small negative effect on host survival (COXME, *A. thailandicus* effect, *p* < 0.001, Fig 1A, B). This deleterious effect was confirmed in an independent genetic background (Oregon R, COXME, *A. thailandicus* effect, p < 0.001, Fig S1). Interestingly, *A. thailandicus* mono-associated flies had 87% lower DCV loads when compared with axenic flies, independently of *Wolbachia* (LMM, *A. thailandicus* effect, *p* < 0.001, Fig 1C). These results show that even though flies with *A. thailandicus* had an increased resistance to DCV, they had a lower tolerance to disease.

**Figure 1.**
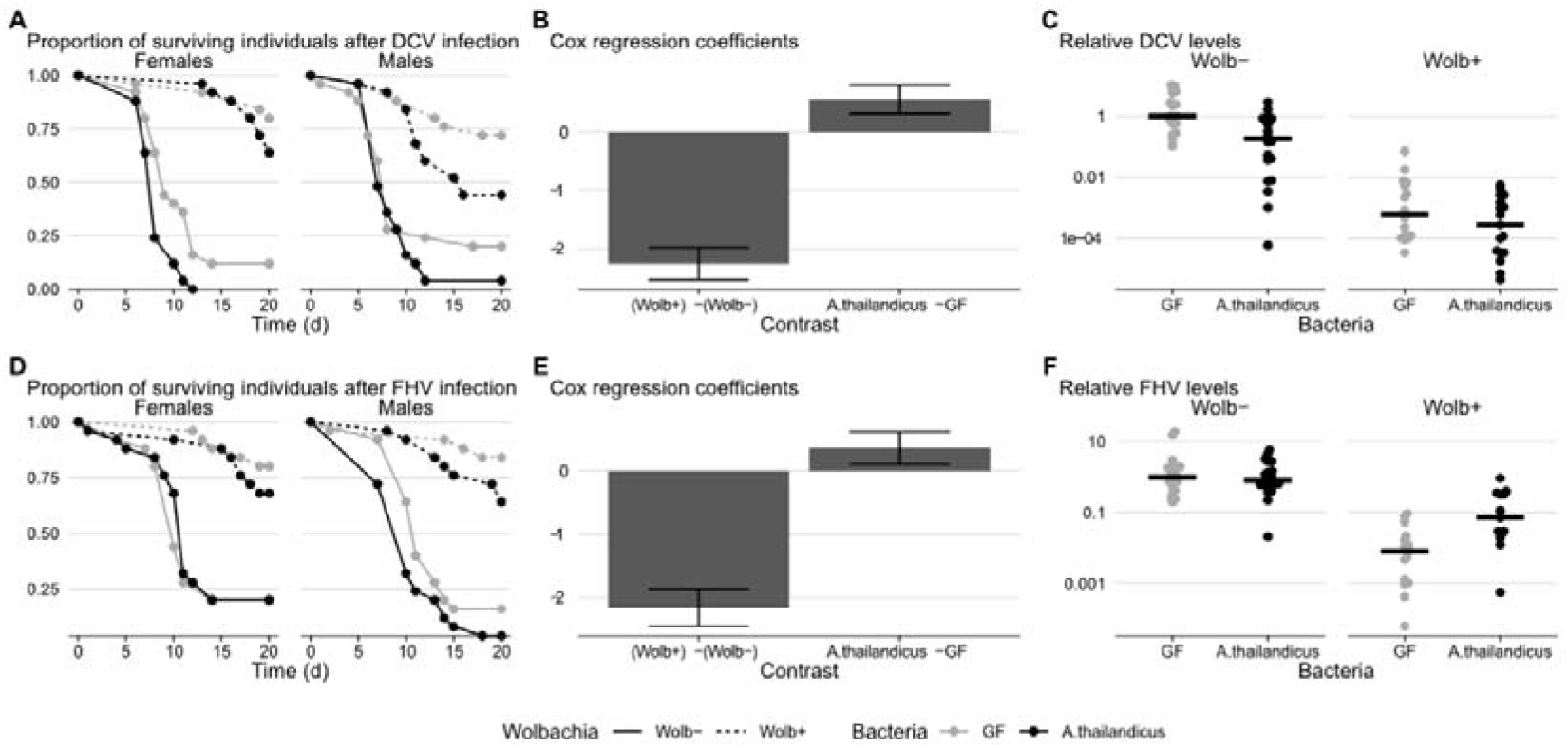
The *Drosophila* gut bacteria A. thailandicus impact viral disease tolerance and resistance. Isogenic *w*^*1118*^ *Drosophila melanogaster* flies with (Wolb+) or without (Wolb-) *Wolbachia*, raised in germ-free conditions (GF) or monoassociated with *A. thailandicus* were systemically infected by pricking with the RNA viruses DCV (10^11^TCID_50_>/ml; A-B) and FHV (10^8^ TCID_50_/ml; D-E) and survival was followed daily for 20 days. Three independent replicates of each sex were performed. Shown are (A,D) the survival curves of each sex for one representative experiment and (B,E) coefficients of the mixed Cox regression model, representing independent effect across all experiments of the presence of *Wolbachia or A. thailandicus* in the mortality risk of flies relative to the mortality risk of Wolbachia-free or gut microbiota free flies. Viral RNA loads of DCV (C) and FHV (F) in infected females were measured in individual flies 48 hours post-infection. Panels represent a single experiment. The experiment was done twice; results were analysed on the aggregate data using mixed effects linear regression models. For (B) and (E) error bars represent upper and lower 95% confidence intervals. For (C) and (F) each dot is a single fly and horizontal bars are medians of the samples.

We repeated these experiments with Flock House Virus (FHV) to test if the effects are conserved when using another RNA virus. Again, we saw no interaction with *Wolbachia* in terms of survival (COXME, *Wolbachia* X A. *thailandicus* effect, p = 0.81; Fig 1D, E) or viral titres (LMM, A. *thailandicus* X *Wolbachia, p* = 0.117, Fig 1F). *Wolbachia* reduced FHV titres 15-fold (LMM, p < 0.001) and increased survival upon infection (COXME, p < 0.001), similarly to what was shown before (Teixeira, Ferreira, and Ashburner 2008; Chrostek et al. 2013). As for DCV, A. *thailandicus* had a small negative effect on survival upon FHV infection (COXME, *p* = 0.006, Fig 1D, E). This negative effect is also observed in Oregon R flies, although in these flies there is an interaction with *Wolbachia* and the effect is stronger in presence of *Wolbachia* (COXME, *A. thailandicus* X *Wolbachia, p* = 0.003, *A. thailandicus* effect *p* < 0.001). However, in contrast to the effect with DCV, FHV loads after infection were 1.83-fold higher in the presence of A. *thailandicus* in the gut of the flies (LMM, p = 0.03, Fig 1D). These results show that this bacterial species led to a decrease in resistance and survival to FHV and variation of the effect of the same gut bacteria on different viruses.

### *Acetobacter thailandicus and Lactobacillus brevis* affect differently host tolerance to systemic viral infection

The previous results showed that the presence of a single bacterial species could have a detrimental impact in disease outcome of systemic viral infection. Other members of the *Drosophila* gut microbiota could impact differently the outcome of the viral infection, either worsening or mitigating the deleterious effects. We therefore tested if *Lactobacillus brevis* also affected DCV infection outcome and if it interacted with *A. thailandicus*. As before, flies associated with *A. thailandicus* showed reduced survival compared with axenic flies (COXME, *A. thailandicus* effect, *p* = 0.002, Fig 2), but there were no changes in survival associated with colonization by *L. brevis*, either alone or in combination with A. thailandicus (COXME, L. brevis effect, p = 0.662, L. brevis * A. *thailandicus effect*, p = 0.155, Fig 2). This shows variation in the effect of different gut colonizing bacteria species on systemic viral infection.

**Figure 2.**
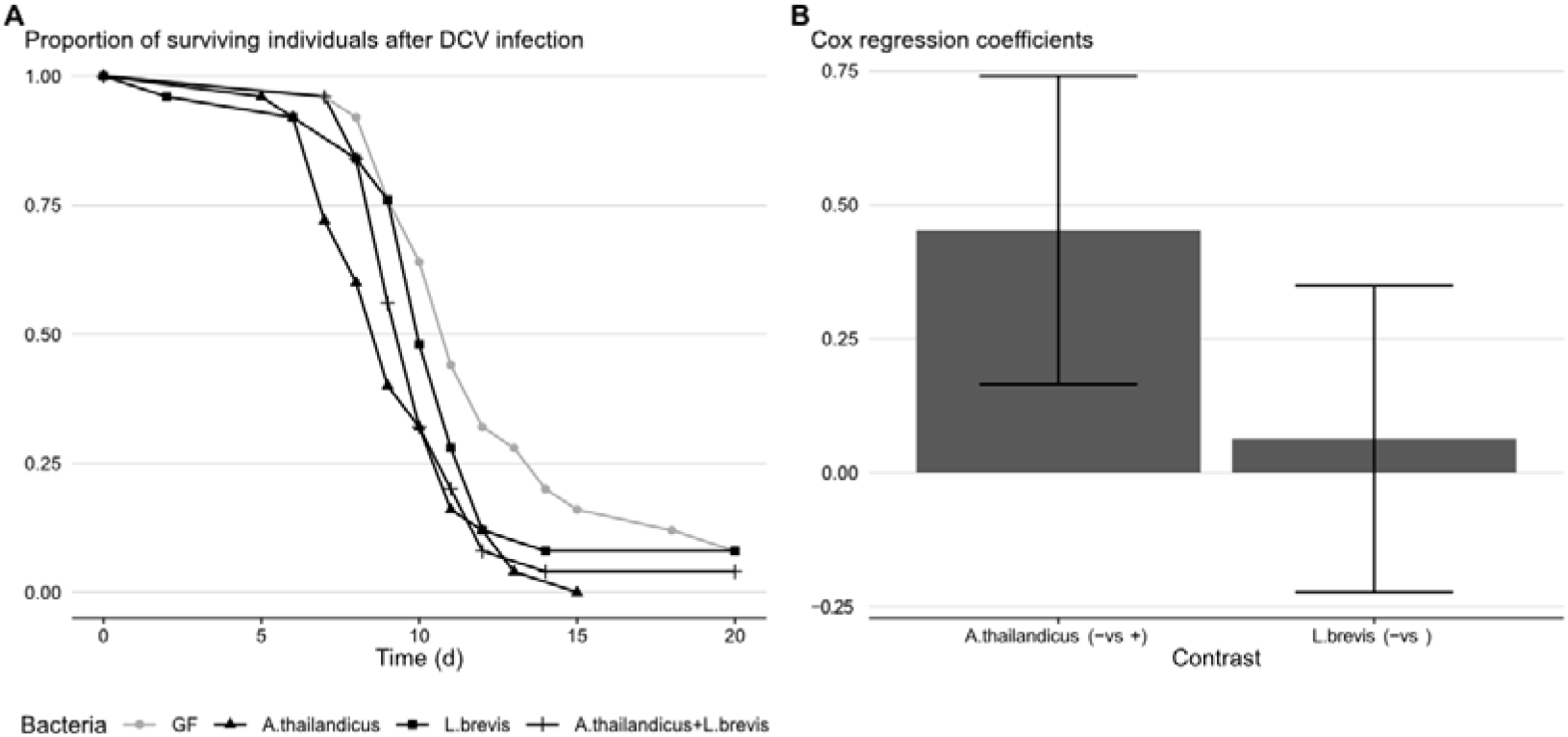
Variation in the effect of Drosophila gut bacteria on DCV infection survival. Isogenic *w*^*1118*^ *Drosophila melanogaster* female flies without *Wolbachia*, in germ-free conditions (GF) or monoassociated with *A. thailandicus, L. brevis* or both bacterial species were systemically infected by pricking with the RNA virus DCV (10^11^ TCID_50_/ml) and survival was followed daily for 20 days. Three independent replicates of were done. Shown are (A) the survival curves for one representative experiment and (B) the coefficients of the mixed Cox regression model, representing the independent effect across all experiments of the presence of *A. thailandicus* or *L. brevis* in the mortality risk of flies relative to the lifespan of flies without that bacteria. For (B) error bars represent upper and lower 95% confidence intervals.

### Presence or absence of gut bacteria do not influence *D. melanogaster disease tolerance* and resistance to oral infection with *Drosophila* C virus

We also tested the effect of the microbiota with DCV oral infection to test if route of infection influences the interaction. Additionally, this route of infection could potentiate the interaction between viruses and bacteria, either directly, or through modulation of the physiology and immune response of the host gut. To test this, we orally infected flies, either raised in axenic conditions or in association with *A. thailandicus, L. brevis* or a mix of both, with DCV and followed survival and measured viral loads two days post infection.

In contrast with the systemic infection route, the presence of *A. thailandicus, L. brevis* or a mix of both did not affect survival (COXME, A. *thailandicus* effect *p* = 0.813, L. brevis effect, *p* = 0.292, *A. thailandicus* * L. brevis effect *p* = 0.519, Fig 3A) or viral loads (Kruskal-Wallis rank sum test, p = 0.818, Fig 3B). These results indicate that the ability of the D. *melanogaster* to tolerate or resist DCV infection was not altered by the presence of these gut bacteria. These results also show that route of viral infection alters the interaction with gut bacteria.

**Figure 3.**
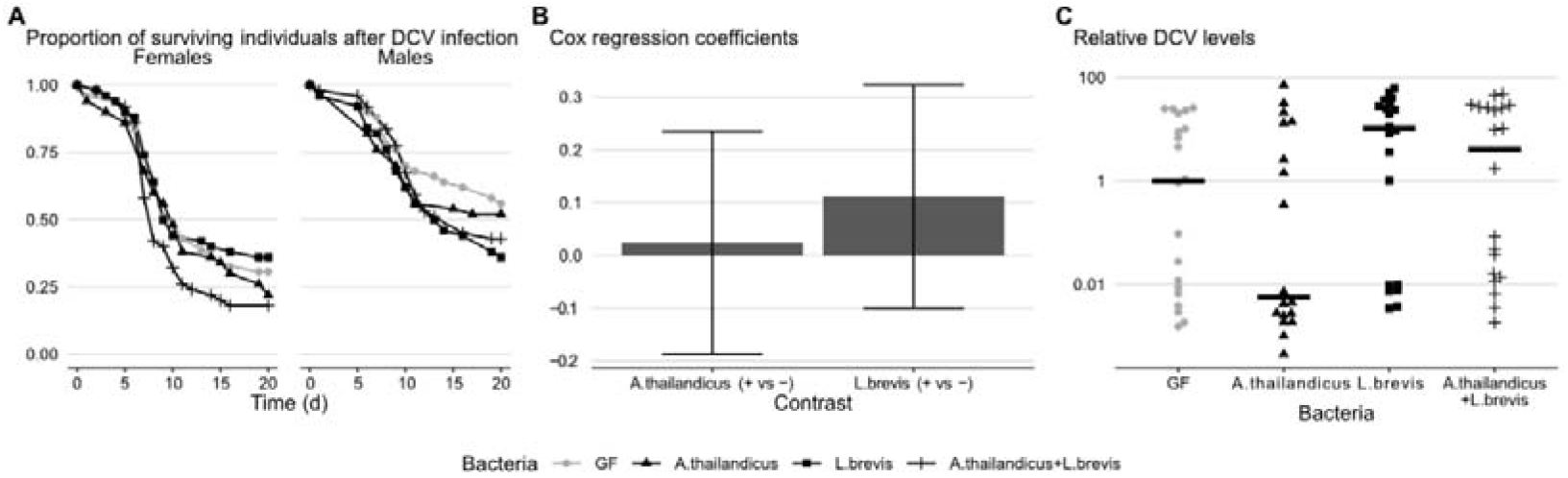
Gut bacteria do not affect host viral resistance and disease tolerance to oral DCV infection. Isogenic *w*^1118^ *Drosophila melanogaster* flies without *Wolbachia*, in germ-free conditions (GF) or monoassociated with *A. thailandicus, L. brevis* or both bacterial species were orally infected with DCV (10^11^ TCID_50_/ml) and survival followed for 20 days. Three independent replicates of each sex were performed. Shown are (A) the survival curves of each sex for one representative experiment and (B) coefficients of the mixed Cox regression model, representing the independent effect across all experiments of the presence of *A. thailandicus* or *L. brevis* in the mortality risk of flies relative to the mortality risk of flies without that bacteria. (C) Viral RNA loads of DCV in infected females were measured in individual flies 48 hours post-infection. Panels represent a single experiment. The experiment was done twice; results were analysed on the aggregate data using a mixed effects linear regression model. For (B) error bars represent upper and lower 95% confidence intervals. For (C) each symbol is a single fly and horizontal bars are medians of the samples.

## Discussion

Our results show that gut-colonizing bacteria associated with *D. melanogaster* in nature can impact host susceptibility to viral infection. However, this effect is contingent on the bacterial species and the infection route. In addition, we show that *Wolbachia*-conferred antiviral protection is not modulated by the presence or absence of these gut bacteria. *Wolbachia*, one of the most prevalent endosymbionts in insects, strongly increases resistance of *D. melanogaster* and other insect to RNA viruses (Hedges et al. 2008; Teixeira, Ferreira, and Ashburner 2008; Moreira et al. 2009). This induction of anti-viral resistance by *Wolbachia* is being deployed as a tool to control mosquito-borne human viral diseases (Moreira et al. 2009; Iturbe-Ormaetxe, Walker, and O Neill 2011), although the molecular mechanisms that explain this viral resistance are still largely unknown. We asked here if gut colonizing bacteria could impact this phenotype. Comparing axenic and mono-associated flies we did not observe an impact on *Wolbachia* mediated anti-viral immunity, confirming previous reports (Ye et al. 2017).

Our work shows that gut-associated bacteria can impact viral infection, and this effect differs with bacterial isolate. *A. thailandicus* mono-association reduces viral titre upon DCV systemic infection but increases lethality of the infection, when compared with axenic flies. Thus, *A. thailandicus* increases resistance to this viral infection, although it reduces disease tolerance. On the other hand, *L. brevis* does not seem to affect either resistance or disease tolerance. This shows that the effect of gut bacteria is species specific, as shown with viral oral infection in *D. melanogaster* and *A. aegypti* (Sansone et al. 2015; Ramirez et al. 2012). The impact of gut bacteria on viral infection also varies with the viruses. Contrary to the effect of *A. thailandicus* on DCV with see an increase in viral titres upon FHV infection. However, *A. thailandicus* decreases D. melanogaster survival to both viruses. Therefore, it is not possible to generalize the effect of these gut bacteria on viral infection in insects.

The comparison of the effect of gut bacteria on viral infection by different routes also shows variation. In contrast to the effect observed on systemic infection, we did not detect any impact of *A. thailandicus* on survival and viral loads after oral DCV infection. Neither did we find an effect of *L. brevis* in this infection. This contrasts with previous reports, where *Acetobacter pomorum* was shown to reduce DCV loads in the gut (Sansone et al. 2015). However, while Sansone et al. measured DCV replication specifically in the gut, we assessed whole body viral levels, a parameter that reflects the action of multiple layers of antiviral protection. Hence, *Acetobacter* species could have a local antiviral role in the gut but not at the whole animal level. It is also possible that different species and isolates of *Acetobacter* have different impact on viral infection.

Overall, the gut-associated bacteria had no or a small impact on resistance and disease tolerance to viruses, particularly when compared with the impact of *Wolbachia*. These results indicate that the effect of gut-associated bacteria on different viral infections routes is intricate, and understanding it requires a detailed characterization of several parameters of infection.

## Supporting information

statistical analysis output

S1_Text Rmd file statistical analysis

## Acknowledgments

This work was funded by Fundação para a Ciência e Tecnologia (FCT) grant PTDC/BEX-GMG/3128/2014 to NM, FCT fellowship FCT-PD/BD/148023/2019 to ML, and FCT grant IF/00839/2015, and ERC grant 773260 to LT.

The fly work at the Fly Facility of Instituto Gulbenkian de Ciencia (Oeiras, Portugal), was partially supported by the research infrastructure Congento, co-financed by Lisboa Regional Operational Programme (Lisboa2020), under the PORTUGAL 2020 Partnership Agreement, through the European Regional Development Fund (ERDF) and Fundação para a Ciência e Tecnologia (Portugal) under the project LISBOA-01-0145-FEDER-022170.

## Supplementary data

**Figure S1.**
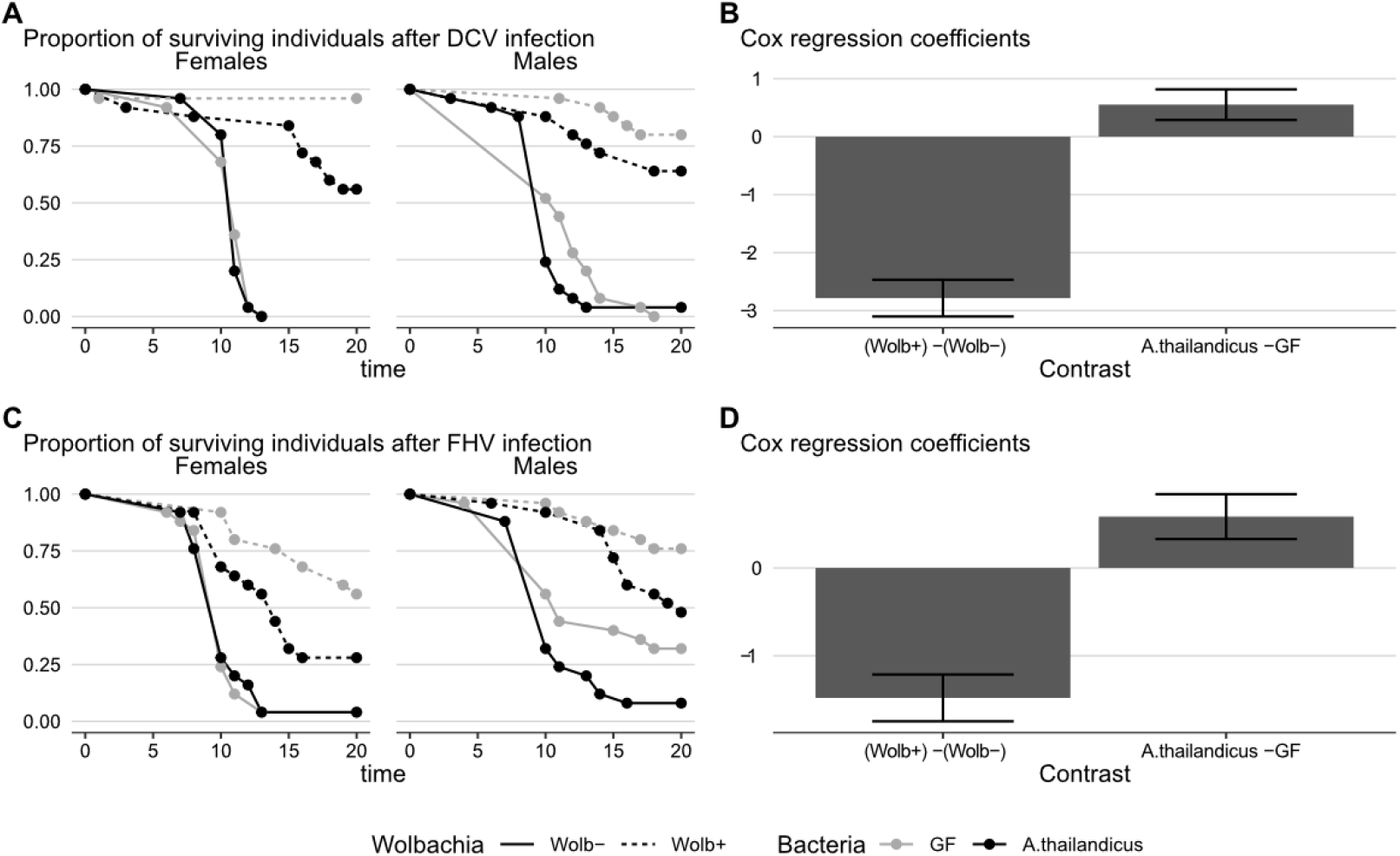
*Drosophila* gut bacteria impact host survival on multiple genetic backgrounds. Oregon R *Drosophila melanogaster* flies with (Wolb+) or without (Wolb-) *Wolbachia*, raised in germ-free conditions (GF) or mono-associated with *A. thailandicus* were systemically infected by pricking with the RNA viruses DCV (10^8^ TCID_50_/ml; A-B) and FHV (10^11^ TCID_50_/ml; C-D) and survival was followed daily for 20 days. Three independent replicates of each sex were performed. Shown are (A, C) the survival curves of each sex for one representative experiment and (B, D) coefficients of the mixed Cox regression model, representing the independent effect across all experiments of the presence of *Wolbachia* or A. *thailandicus* in the mortality risk of flies relative to the mortality risk of *Wolbachia*-or gut microbiota free flies. For (B) and (D) error bars represent upper and lower confidence intervals.

**S1 Text** – Rmd script with statistical analysis

**S1 Data** – csv file with survival data

**S2 Data** – csv file with qPCR viral titre data

