## Supplementary material for "Viral route of infection determines the effect of *Drosophila melanogaster* gut bacteria on host resistance and tolerance to disease": statistical analysis output: Microbiota_and_viruses_Analysis.html

Microbiota and Viral Infection


### Microbiota and Viral Infection

#### Data input

##### Survival

```
surv=read_csv(here("data/DatasetS1_survival.csv")) %>% 
  mutate(Wolbachia=fct_relevel(Wolbachia,"Wolb-"),
         Stock=fct_relevel(Stock,"W1118"),
         Bacteria=fct_relevel(Bacteria,
                              "GF","A.thailandicus","L.brevis")
         )
```

```
## 
## -- Column specification --------------------------------------------------------
## cols(
##   Experiment.type = col_character(),
##   Virus = col_character(),
##   Infection = col_character(),
##   Stock = col_character(),
##   Wolbachia = col_character(),
##   Bacteria = col_character(),
##   Sex = col_character(),
##   Experiment = col_character(),
##   Replicate = col_double(),
##   RepFull = col_character(),
##   day_status = col_double(),
##   status = col_double()
## )
```

###### Systemic infection, colonization by A.thailandicus

```
surv_systemic=surv %>% 
  filter(Infection=="Systemic",`Experiment.type`=="Microbiota_and_Wolbachia") %>% 
  droplevels()
  
surv_systemic %>% 
  group_by(Virus,Infection,Stock,Wolbachia,Bacteria,Sex,Experiment) %>% 
    tally
```

```
## # A tibble: 160 x 8
## # Groups:   Virus, Infection, Stock, Wolbachia, Bacteria, Sex [64]
##    Virus Infection Stock Wolbachia Bacteria       Sex     Experiment     n
##    <chr> <chr>     <fct> <fct>     <fct>          <chr>   <chr>      <int>
##  1 DCV   Systemic  W1118 Wolb-     GF             Females A             25
##  2 DCV   Systemic  W1118 Wolb-     GF             Females B             25
##  3 DCV   Systemic  W1118 Wolb-     GF             Females C             25
##  4 DCV   Systemic  W1118 Wolb-     GF             Males   A             25
##  5 DCV   Systemic  W1118 Wolb-     GF             Males   B             25
##  6 DCV   Systemic  W1118 Wolb-     GF             Males   C             25
##  7 DCV   Systemic  W1118 Wolb-     A.thailandicus Females A             25
##  8 DCV   Systemic  W1118 Wolb-     A.thailandicus Females B             25
##  9 DCV   Systemic  W1118 Wolb-     A.thailandicus Females C             25
## 10 DCV   Systemic  W1118 Wolb-     A.thailandicus Males   A             25
## # ... with 150 more rows
```

###### Systemic infection, colonization by A.thailandicus and L. brevis

```
surv_systemic_multi=surv %>% 
  filter(Infection=="Systemic",`Experiment.type`=="Different_microbiota") %>% 
  droplevels()
  
surv_systemic_multi %>% 
  group_by(Virus,Bacteria,Sex,Experiment) %>% 
    tally
```

```
## # A tibble: 20 x 5
## # Groups:   Virus, Bacteria, Sex [8]
##    Virus Bacteria                Sex     Experiment     n
##    <chr> <fct>                   <chr>   <chr>      <int>
##  1 DCV   GF                      Females A             25
##  2 DCV   GF                      Females B             25
##  3 DCV   GF                      Females C             25
##  4 DCV   A.thailandicus          Females A             25
##  5 DCV   A.thailandicus          Females B             25
##  6 DCV   A.thailandicus          Females C             25
##  7 DCV   L.brevis                Females A             25
##  8 DCV   L.brevis                Females B             25
##  9 DCV   L.brevis                Females C             25
## 10 DCV   A.thailandicus+L.brevis Females A             25
## 11 DCV   A.thailandicus+L.brevis Females B             25
## 12 DCV   A.thailandicus+L.brevis Females C             25
## 13 Mock  GF                      Females A             25
## 14 Mock  GF                      Females B             25
## 15 Mock  A.thailandicus          Females A             25
## 16 Mock  A.thailandicus          Females B             25
## 17 Mock  L.brevis                Females A             25
## 18 Mock  L.brevis                Females B             25
## 19 Mock  A.thailandicus+L.brevis Females A             25
## 20 Mock  A.thailandicus+L.brevis Females B             25
```

###### Oral infection by A.thailandicus and L. brevis

```
surv_oral=surv %>% 
  filter(Infection=="Oral") %>% 
  droplevels()
  
surv_oral %>% 
    group_by(Virus,Stock,Bacteria,Sex,Experiment) %>% 
  tally
```

```
## # A tibble: 48 x 6
## # Groups:   Virus, Stock, Bacteria, Sex [16]
##    Virus Stock Bacteria       Sex     Experiment     n
##    <chr> <fct> <fct>          <chr>   <chr>      <int>
##  1 DCV   W1118 GF             Females A             44
##  2 DCV   W1118 GF             Females B             48
##  3 DCV   W1118 GF             Females C             49
##  4 DCV   W1118 GF             Males   A             50
##  5 DCV   W1118 GF             Males   B             50
##  6 DCV   W1118 GF             Males   C             50
##  7 DCV   W1118 A.thailandicus Females A             46
##  8 DCV   W1118 A.thailandicus Females B             49
##  9 DCV   W1118 A.thailandicus Females C             50
## 10 DCV   W1118 A.thailandicus Males   A             50
## # ... with 38 more rows
```

##### Loads

```
viral_levels=read.csv(here("data/DatasetS2_viral_loads.csv")) %>% 
  mutate(Bacteria=fct_relevel(Bacteria,
                              "GF",
                              "A.thailandicus",
                              "L.brevis"))
```

###### Systemic infection by DCV

```
levels_systemic_DCV= viral_levels %>% 
  filter(Virus=="DCV",Infection=="Systemic") %>% 
  droplevels()

head(levels_systemic_DCV)
```

```
##   Infection Virus Wolbachia       Bacteria Replicate fold_change
## 1  Systemic   DCV     Wolb- A.thailandicus         A 0.307121046
## 2  Systemic   DCV     Wolb- A.thailandicus         A 2.481279723
## 3  Systemic   DCV     Wolb- A.thailandicus         A 0.000403169
## 4  Systemic   DCV     Wolb- A.thailandicus         A 0.004413968
## 5  Systemic   DCV     Wolb- A.thailandicus         A 0.248400930
## 6  Systemic   DCV     Wolb- A.thailandicus         A 0.000020000
##   normalized_fold_change
## 1            0.367037071
## 2            2.965350808
## 3            0.000481823
## 4            0.005275086
## 5            0.296861289
## 6            0.000023900
```

###### Systemic infection by FHV

```
levels_systemic_FHV= viral_levels %>% 
  filter(Virus=="FHV",Infection=="Systemic") %>% 
  droplevels()

head(levels_systemic_FHV)
```

```
##   Infection Virus Wolbachia       Bacteria Replicate fold_change
## 1  Systemic   FHV     Wolb- A.thailandicus         A  6.60028276
## 2  Systemic   FHV     Wolb- A.thailandicus         A  1.70401308
## 3  Systemic   FHV     Wolb- A.thailandicus         A  2.07759429
## 4  Systemic   FHV     Wolb- A.thailandicus         A  0.92045395
## 5  Systemic   FHV     Wolb- A.thailandicus         A  0.02961264
## 6  Systemic   FHV     Wolb- A.thailandicus         A  0.83563460
##   normalized_fold_change
## 1             4.65918003
## 2             1.20287327
## 3             1.46658653
## 4             0.64975408
## 5             0.02090375
## 6             0.58987958
```

###### Oral infection by DCV

```
levels_oral_DCV= viral_levels %>% 
  filter(Virus=="DCV",Infection=="Oral") %>% 
  droplevels()

head(levels_oral_DCV)
```

```
##   Infection Virus Wolbachia       Bacteria Replicate  fold_change
## 1      Oral   DCV     Wolb- A.thailandicus         A 4.095451e-03
## 2      Oral   DCV     Wolb- A.thailandicus         A 1.937841e+02
## 3      Oral   DCV     Wolb- A.thailandicus         A 2.876064e+02
## 4      Oral   DCV     Wolb- A.thailandicus         A 2.247753e-02
## 5      Oral   DCV     Wolb- A.thailandicus         A 2.506705e-02
## 6      Oral   DCV     Wolb- A.thailandicus         A 6.545664e-02
##   normalized_fold_change
## 1            0.000449157
## 2           21.252710370
## 3           31.542403180
## 4            0.002465158
## 5            0.002749157
## 6            0.007178767
```

#### Figure 1 - Systemic infection, w1118

##### Survival

###### DCV

###### Data selection

```
surv_systemic_DCV=filter(surv_systemic,
                         Virus=="DCV",
                         Stock=="W1118")
```

###### Mixed Effects Cox Model

- Full model
- Effect of Wolbachia, Bacteria and Sex, but no interaction

```
surv_systemic_DCV_mixed_full=coxme(Surv(day_status,status)~
                                     Wolbachia*Bacteria*Sex
                                   +(1|Experiment)+(1|RepFull),
                                   surv_systemic_DCV)

summary(surv_systemic_DCV_mixed_full)
```

```
## Cox mixed-effects model fit by maximum likelihood
##   Data: surv_systemic_DCV
##   events, n = 405, 600
##   Iterations= 10 63 
##                     NULL Integrated    Fitted
## Log-likelihood -2405.485  -2213.097 -2173.355
## 
##                    Chisq    df p    AIC    BIC
## Integrated loglik 384.77  9.00 0 366.77 330.74
##  Penalized loglik 464.26 40.38 0 383.49 221.80
## 
## Model:  Surv(day_status, status) ~ Wolbachia * Bacteria * Sex + (1 |      Experiment) + (1 | RepFull) 
## Fixed coefficients
##                                                       coef exp(coef)  se(coef)
## WolbachiaWolb+                                 -2.20325257 0.1104433 0.2875547
## BacteriaA.thailandicus                          0.60658743 1.8341615 0.2181918
## SexMales                                        0.52797492 1.6954953 0.2216456
## WolbachiaWolb+:BacteriaA.thailandicus          -0.06435006 0.9376767 0.3766125
## WolbachiaWolb+:SexMales                        -0.08637818 0.9172473 0.3897064
## BacteriaA.thailandicus:SexMales                -0.07548710 0.9272917 0.3081223
## WolbachiaWolb+:BacteriaA.thailandicus:SexMales  0.08158862 1.0850094 0.5187299
##                                                    z       p
## WolbachiaWolb+                                 -7.66 1.8e-14
## BacteriaA.thailandicus                          2.78 5.4e-03
## SexMales                                        2.38 1.7e-02
## WolbachiaWolb+:BacteriaA.thailandicus          -0.17 8.6e-01
## WolbachiaWolb+:SexMales                        -0.22 8.2e-01
## BacteriaA.thailandicus:SexMales                -0.24 8.1e-01
## WolbachiaWolb+:BacteriaA.thailandicus:SexMales  0.16 8.8e-01
## 
## Random effects
##  Group      Variable  Std Dev     Variance   
##  Experiment Intercept 0.099956210 0.009991244
##  RepFull    Intercept 0.364878968 0.133136661
```

```
Anova(surv_systemic_DCV_mixed_full)
```

```
## Analysis of Deviance Table (Type II tests)
## 
## Response: Surv(day_status, status)
##                        Df    Chisq Pr(>Chisq)    
## Wolbachia               1 263.8569  < 2.2e-16 ***
## Bacteria                1  20.2233  6.891e-06 ***
## Sex                     1  14.6261  0.0001311 ***
## Wolbachia:Bacteria      1   0.0068  0.9341564    
## Wolbachia:Sex           1   0.0246  0.8754488    
## Bacteria:Sex            1   0.0355  0.8505614    
## Wolbachia:Bacteria:Sex  1   0.0247  0.8750199    
## ---
## Signif. codes:  0 '***' 0.001 '**' 0.01 '*' 0.05 '.' 0.1 ' ' 1
```

- Simplified model

```
surv_systemic_DCV_mixed_reduced=coxme(Surv(day_status,status)~
                                           Wolbachia+Bacteria+Sex
                                      +(1|Experiment)+(1|RepFull),
                                         surv_systemic_DCV)

summary(surv_systemic_DCV_mixed_reduced)
```

```
## Cox mixed-effects model fit by maximum likelihood
##   Data: surv_systemic_DCV
##   events, n = 405, 600
##   Iterations= 10 53 
##                     NULL Integrated    Fitted
## Log-likelihood -2405.485  -2213.143 -2173.842
## 
##                    Chisq    df p    AIC    BIC
## Integrated loglik 384.68  5.00 0 374.68 354.66
##  Penalized loglik 463.28 37.18 0 388.92 240.04
## 
## Model:  Surv(day_status, status) ~ Wolbachia + Bacteria + Sex + (1 |      Experiment) + (1 | RepFull) 
## Fixed coefficients
##                              coef exp(coef)  se(coef)      z       p
## WolbachiaWolb+         -2.2586475 0.1044917 0.1389681 -16.25 0.0e+00
## BacteriaA.thailandicus  0.5611107 1.7526180 0.1244272   4.51 6.5e-06
## SexMales                0.4728683 1.6045900 0.1234888   3.83 1.3e-04
## 
## Random effects
##  Group      Variable  Std Dev     Variance   
##  Experiment Intercept 0.099930373 0.009986079
##  RepFull    Intercept 0.362178856 0.131173524
```

```
Anova(surv_systemic_DCV_mixed_reduced)
```

```
## Analysis of Deviance Table (Type II tests)
## 
## Response: Surv(day_status, status)
##           Df   Chisq Pr(>Chisq)    
## Wolbachia  1 264.160  < 2.2e-16 ***
## Bacteria   1  20.336  6.496e-06 ***
## Sex        1  14.663  0.0001285 ***
## ---
## Signif. codes:  0 '***' 0.001 '**' 0.01 '*' 0.05 '.' 0.1 ' ' 1
```

- Average effects of *Wolbachia* and *A. thailandicus* presence
- Decreased risk in the presence of Wolbachia and increased risk in the presence of *A. thailandicus*

```
surv_systemic_DCV_mixed_emmeans=emmeans(surv_systemic_DCV_mixed_reduced,
                                        list(Bacteria=trt.vs.ctrl~Bacteria,
                                             wolb=trt.vs.ctrl~Wolbachia))

surv_systemic_DCV_mixed_emmeans
```

```
## $`Bacteria emmeans`
##  Bacteria       emmean     SE  df asymp.LCL asymp.UCL
##  GF             -0.281 0.0622 Inf    -0.402    -0.159
##  A.thailandicus  0.281 0.0622 Inf     0.159     0.402
## 
## Results are averaged over the levels of: Wolbachia, Sex 
## Results are given on the log (not the response) scale. 
## Confidence level used: 0.95 
## 
## $`Bacteria differences from control`
##  Bacteria.contrast   estimate    SE  df z.ratio p.value
##  A.thailandicus - GF    0.561 0.124 Inf 4.510   <.0001 
## 
## Results are averaged over the levels of: Wolbachia, Sex 
## Results are given on the log (not the response) scale. 
## 
## $`wolb emmeans`
##  Wolbachia emmean     SE  df asymp.LCL asymp.UCL
##  Wolb-       1.13 0.0695 Inf     0.993     1.266
##  Wolb+      -1.13 0.0695 Inf    -1.266    -0.993
## 
## Results are averaged over the levels of: Bacteria, Sex 
## Results are given on the log (not the response) scale. 
## Confidence level used: 0.95 
## 
## $`wolb differences from control`
##  wolb.contrast     estimate    SE  df z.ratio p.value
##  (Wolb+) - (Wolb-)    -2.26 0.139 Inf -16.253 <.0001 
## 
## Results are averaged over the levels of: Bacteria, Sex 
## Results are given on the log (not the response) scale.
```

###### Plots

- Overall plot of all experiments

```
surv_systemic_DCV_fit=survfit(Surv(day_status,status)~Bacteria+Wolbachia+Experiment+Sex,surv_systemic_DCV)

surv_systemic_DCV_summary=surv_summary(surv_systemic_DCV_fit,data=surv_systemic_DCV) %>% 
  mutate(Bacteria=fct_relevel(Bacteria,"GF")) %>%
  group_by(strata,Bacteria,Sex,Wolbachia,Experiment) %>% 
  group_modify(~{
    if(min(.x$time)!=0){.x=add_row(.x,time=0,surv=1)}else{.x}})

surv_systemic_DCV_plot=ggplot(surv_systemic_DCV_summary)+
  aes(x=time,y=surv)+
  geom_line(aes(linetype=Wolbachia,color=Bacteria))+
  geom_point(aes(color=Bacteria))+
  scale_color_manual(values=c("darkgray","black"))+
  scale_y_continuous(limits=c(0,1))+
  facet_grid(Sex~Experiment)+
  labs(y=NULL,x="Time (d)")+
  theme_HMI()
  
surv_systemic_DCV_plot
```

- Panels chosen for figure

```
surv_systemic_DCV_panel=surv_systemic_DCV_plot%+%
  filter(surv_systemic_DCV_summary,Experiment=="C")+
  facet_grid(~Sex)+
  scale_y_continuous(expand = c(0,0))+
  labs(title="Proportion of surviving individuals after DCV infection")
```

```
## Scale for 'y' is already present. Adding another scale for 'y', which will
## replace the existing scale.
```

```
surv_systemic_DCV_panel
```

- Hazard ratio plot

```
surv_systemic_DCV_hazards=map_dfr(c(2,4),~
                                    surv_systemic_DCV_mixed_emmeans[[.x]] %>% confint%>% tidy) %>% 
  mutate(contrast=coalesce(wolb.contrast,Bacteria.contrast))

surv_systemic_DCV_hazards
```

```
## # A tibble: 2 x 8
##   Bacteria.contra~ estimate std.error    df asymp.LCL asymp.UCL wolb.contrast
##   <chr>               <dbl>     <dbl> <dbl>     <dbl>     <dbl> <chr>        
## 1 A.thailandicus ~    0.561     0.124   Inf     0.317     0.805 <NA>         
## 2 <NA>               -2.26      0.139   Inf    -2.53     -1.99  (Wolb+) - (W~
## # ... with 1 more variable: contrast <chr>
```

```
surv_systemic_DCV_hazards_plot=ggplot(surv_systemic_DCV_hazards)+
  aes(x=contrast,y=estimate)+
  geom_bar(aes(x=contrast,y=estimate),stat="identity")+
  geom_errorbar(aes(ymin=asymp.LCL,ymax=asymp.UCL),width=.5)+
  facet_grid(~df)+
  theme_HMI() +
  labs(title="Cox regression coefficients",y=NULL,x="Contrast")


surv_systemic_DCV_hazards_plot
```

###### FHV

###### Data selection

```
surv_systemic_FHV=filter(surv_systemic,Virus%in%c("FHV"),Stock=="W1118") %>% 
  mutate(Wolbachia=fct_relevel(Wolbachia,"Wolb-"),
         Bacteria=fct_relevel(Bacteria,"GF"))
```

###### Mixed effect model

- Full model

  - Effect of Wolbachia, Bacteria and Sex, but no interaction

```
surv_systemic_FHV_model_mixed_full=coxme(Surv(day_status,status)~
                                           Wolbachia*Bacteria*Sex+
                                           (1|Experiment)+
                                           (1|RepFull),
                                         surv_systemic_FHV)


Anova(surv_systemic_FHV_model_mixed_full)
```

```
## Analysis of Deviance Table (Type II tests)
## 
## Response: Surv(day_status, status)
##                        Df    Chisq Pr(>Chisq)    
## Wolbachia               1 202.6431  < 2.2e-16 ***
## Bacteria                1   7.6168   0.005783 ** 
## Sex                     1   5.3400   0.020841 *  
## Wolbachia:Bacteria      1   0.0583   0.809175    
## Wolbachia:Sex           1   2.4996   0.113873    
## Bacteria:Sex            1   0.1140   0.735615    
## Wolbachia:Bacteria:Sex  1   0.9812   0.321892    
## ---
## Signif. codes:  0 '***' 0.001 '**' 0.01 '*' 0.05 '.' 0.1 ' ' 1
```

```
summary(surv_systemic_FHV_model_mixed_full)
```

```
## Cox mixed-effects model fit by maximum likelihood
##   Data: surv_systemic_FHV
##   events, n = 343, 600
##   Iterations= 24 147 
##                     NULL Integrated    Fitted
## Log-likelihood -2069.469  -1913.408 -1881.679
## 
##                    Chisq    df p    AIC    BIC
## Integrated loglik 312.12  9.00 0 294.12 259.58
##  Penalized loglik 375.58 34.14 0 307.29 176.26
## 
## Model:  Surv(day_status, status) ~ Wolbachia * Bacteria * Sex + (1 |      Experiment) + (1 | RepFull) 
## Fixed coefficients
##                                                      coef exp(coef)  se(coef)
## WolbachiaWolb+                                 -2.0468511 0.1291409 0.2862655
## BacteriaA.thailandicus                          0.2596449 1.2964697 0.2195863
## SexMales                                       -0.3049049 0.7371935 0.2222928
## WolbachiaWolb+:BacteriaA.thailandicus           0.1742586 1.1903634 0.3818402
## WolbachiaWolb+:SexMales                        -0.1548593 0.8565357 0.4302642
## BacteriaA.thailandicus:SexMales                 0.2496740 1.2836069 0.3093962
## WolbachiaWolb+:BacteriaA.thailandicus:SexMales -0.5844210 0.5574285 0.5899804
##                                                    z       p
## WolbachiaWolb+                                 -7.15 8.7e-13
## BacteriaA.thailandicus                          1.18 2.4e-01
## SexMales                                       -1.37 1.7e-01
## WolbachiaWolb+:BacteriaA.thailandicus           0.46 6.5e-01
## WolbachiaWolb+:SexMales                        -0.36 7.2e-01
## BacteriaA.thailandicus:SexMales                 0.81 4.2e-01
## WolbachiaWolb+:BacteriaA.thailandicus:SexMales -0.99 3.2e-01
## 
## Random effects
##  Group      Variable  Std Dev      Variance    
##  Experiment Intercept 0.0132340246 0.0001751394
##  RepFull    Intercept 0.3521875568 0.1240360752
```

- Simplified model

```
surv_systemic_FHV_model_mixed_reduced=coxme(Surv(day_status,status)~
                                          Wolbachia+Bacteria+Sex+Experiment
                                          +(1|RepFull),
                                         surv_systemic_FHV)


Anova(surv_systemic_FHV_model_mixed_reduced)
```

```
## Analysis of Deviance Table (Type II tests)
## 
## Response: Surv(day_status, status)
##            Df    Chisq Pr(>Chisq)    
## Wolbachia   1 209.8623  < 2.2e-16 ***
## Bacteria    1   7.4407   0.006377 ** 
## Sex         1   5.6776   0.017182 *  
## Experiment  2   1.3512   0.508861    
## ---
## Signif. codes:  0 '***' 0.001 '**' 0.01 '*' 0.05 '.' 0.1 ' ' 1
```

```
summary(surv_systemic_FHV_model_mixed_reduced)
```

```
## Cox mixed-effects model fit by maximum likelihood
##   Data: surv_systemic_FHV
##   events, n = 343, 600
##   Iterations= 10 54 
##                     NULL Integrated    Fitted
## Log-likelihood -2069.469  -1914.611 -1881.132
## 
##                    Chisq    df p    AIC    BIC
## Integrated loglik 309.72  6.00 0 297.72 274.69
##  Penalized loglik 376.67 33.88 0 308.91 178.87
## 
## Model:  Surv(day_status, status) ~ Wolbachia + Bacteria + Sex + Experiment +      (1 | RepFull) 
## Fixed coefficients
##                               coef exp(coef)  se(coef)      z      p
## WolbachiaWolb+         -2.16158116 0.1151429 0.1492122 -14.49 0.0000
## BacteriaA.thailandicus  0.36131210 1.4352113 0.1324575   2.73 0.0064
## SexMales               -0.31487456 0.7298804 0.1321458  -2.38 0.0170
## ExperimentB            -0.08337181 0.9200090 0.1606798  -0.52 0.6000
## ExperimentC            -0.18779747 0.8287825 0.1618061  -1.16 0.2500
## 
## Random effects
##  Group   Variable  Std Dev   Variance 
##  RepFull Intercept 0.3637777 0.1323342
```

- Average effect of *Wolbachia* and *A. thailandicus* presence
- Decreased risk in the presence of Wolbachia and increased risk in the presence of *A. thailandicus*

```
surv_systemic_FHV_mixed_emmeans=emmeans(surv_systemic_FHV_model_mixed_reduced,
                                        list(Bacteria=trt.vs.ctrl~Bacteria,
                                             wolb=trt.vs.ctrl~Wolbachia))

surv_systemic_FHV_mixed_emmeans
```

```
## $`Bacteria emmeans`
##  Bacteria       emmean     SE  df asymp.LCL asymp.UCL
##  GF             -0.181 0.0662 Inf   -0.3105   -0.0509
##  A.thailandicus  0.181 0.0662 Inf    0.0509    0.3105
## 
## Results are averaged over the levels of: Wolbachia, Sex, Experiment 
## Results are given on the log (not the response) scale. 
## Confidence level used: 0.95 
## 
## $`Bacteria differences from control`
##  Bacteria.contrast   estimate    SE  df z.ratio p.value
##  A.thailandicus - GF    0.361 0.132 Inf 2.728   0.0064 
## 
## Results are averaged over the levels of: Wolbachia, Sex, Experiment 
## Results are given on the log (not the response) scale. 
## 
## $`wolb emmeans`
##  Wolbachia emmean     SE  df asymp.LCL asymp.UCL
##  Wolb-       1.08 0.0746 Inf     0.935     1.227
##  Wolb+      -1.08 0.0746 Inf    -1.227    -0.935
## 
## Results are averaged over the levels of: Bacteria, Sex, Experiment 
## Results are given on the log (not the response) scale. 
## Confidence level used: 0.95 
## 
## $`wolb differences from control`
##  wolb.contrast     estimate    SE  df z.ratio p.value
##  (Wolb+) - (Wolb-)    -2.16 0.149 Inf -14.487 <.0001 
## 
## Results are averaged over the levels of: Bacteria, Sex, Experiment 
## Results are given on the log (not the response) scale.
```

###### Plots

- Overall plot of all experiments

```
surv_systemic_FHV_fit=survfit(Surv(day_status,status)~Bacteria+Wolbachia+Experiment+Sex,surv_systemic_FHV)

surv_systemic_FHV_summary=surv_summary(surv_systemic_FHV_fit,data=surv_systemic_FHV) %>% 
  mutate(Bacteria=fct_relevel(Bacteria,"GF")) %>% 
  group_by(strata,Bacteria,Wolbachia,Sex,Experiment) %>% 
  group_modify(~{
    if(min(.x$time)!=0){.x=add_row(.x,time=0,surv=1)}else{.x}})

surv_systemic_FHV_plot=ggplot(surv_systemic_FHV_summary)+
  aes(x=time,y=surv)+
  geom_line(aes(linetype=Wolbachia,color=Bacteria))+
  geom_point(aes(color=Bacteria))+
  scale_color_manual(values=c("darkgray","black"))+
  scale_y_continuous(limits=c(0,1))+
  facet_grid(Sex~Experiment)+
  labs(y="Survival",x="Time (d)")+
  theme_HMI()
  
surv_systemic_FHV_plot
```

- Panels chosen for figure

```
surv_systemic_FHV_panel=surv_systemic_FHV_plot%+%
  filter(surv_systemic_FHV_summary,Experiment=="C")+
  facet_grid(~Sex)+
  scale_y_continuous(expand = c(0,0))+
  labs(title="Proportion of surviving individuals after FHV infection",y=NULL)
```

```
## Scale for 'y' is already present. Adding another scale for 'y', which will
## replace the existing scale.
```

```
surv_systemic_FHV_panel
```

- Hazard ratio plots

```
surv_systemic_FHV_hazards=map_dfr(c(2,4),~
                                    surv_systemic_FHV_mixed_emmeans[[.x]] %>% confint%>% tidy) %>% 
  mutate(contrast=coalesce(wolb.contrast,Bacteria.contrast))

surv_systemic_FHV_hazards
```

```
## # A tibble: 2 x 8
##   Bacteria.contra~ estimate std.error    df asymp.LCL asymp.UCL wolb.contrast
##   <chr>               <dbl>     <dbl> <dbl>     <dbl>     <dbl> <chr>        
## 1 A.thailandicus ~    0.361     0.132   Inf     0.102     0.621 <NA>         
## 2 <NA>               -2.16      0.149   Inf    -2.45     -1.87  (Wolb+) - (W~
## # ... with 1 more variable: contrast <chr>
```

```
surv_systemic_FHV_hazards_plot=ggplot(surv_systemic_FHV_hazards)+
  aes(x=contrast,y=estimate)+
  geom_bar(aes(x=contrast,y=estimate),stat="identity")+
  geom_errorbar(aes(ymin=asymp.LCL,ymax=asymp.UCL),width=.5)+
  theme_HMI() +
  labs(title="Cox regression coefficients",y=NULL,x="Contrast")


surv_systemic_FHV_hazards_plot
```

##### Virus Levels

###### DCV

###### Mixed effect model

- Full model

  - No interaction between microbiota and wolbachia, but a significant reduction of viral loads in the presence of *A.thailandicus*

```
levels_systemic_DCV_model_full=lmer(log10(normalized_fold_change)~
                                 Bacteria*Wolbachia+(1|Replicate),
                               data=levels_systemic_DCV)


Anova(levels_systemic_DCV_model_full)
```

```
## Analysis of Deviance Table (Type II Wald chisquare tests)
## 
## Response: log10(normalized_fold_change)
##                       Chisq Df Pr(>Chisq)    
## Bacteria            41.5430  1  1.153e-10 ***
## Wolbachia          573.4358  1  < 2.2e-16 ***
## Bacteria:Wolbachia   2.8284  1    0.09261 .  
## ---
## Signif. codes:  0 '***' 0.001 '**' 0.01 '*' 0.05 '.' 0.1 ' ' 1
```

```
summary(levels_systemic_DCV_model_full)
```

```
## Linear mixed model fit by REML ['lmerMod']
## Formula: log10(normalized_fold_change) ~ Bacteria * Wolbachia + (1 | Replicate)
##    Data: levels_systemic_DCV
## 
## REML criterion at convergence: 564.6
## 
## Scaled residuals: 
##     Min      1Q  Median      3Q     Max 
## -3.5358 -0.5981  0.1029  0.7166  2.6169 
## 
## Random effects:
##  Groups    Name        Variance Std.Dev.
##  Replicate (Intercept) 0.04666  0.2160  
##  Residual              0.89767  0.9475  
## Number of obs: 204, groups:  Replicate, 3
## 
## Fixed effects:
##                                       Estimate Std. Error t value
## (Intercept)                           -0.02135    0.17772  -0.120
## BacteriaA.thailandicus                -1.06312    0.17605  -6.039
## WolbachiaWolb+                        -3.46391    0.18071 -19.168
## BacteriaA.thailandicus:WolbachiaWolb+  0.46400    0.27590   1.682
## 
## Correlation of Fixed Effects:
##             (Intr) BctrA. WlbcW+
## BctrA.thlnd -0.512              
## WolbachWlb+ -0.499  0.504       
## BctrA.t:WW+  0.326 -0.637 -0.656
```

- Simplified model

```
levels_systemic_DCV_model_reduced=lmer(log10(normalized_fold_change)~
                                         Bacteria+Wolbachia+(1|Replicate),
                               data=levels_systemic_DCV)
Anova(levels_systemic_DCV_model_reduced)
```

```
## Analysis of Deviance Table (Type II Wald chisquare tests)
## 
## Response: log10(normalized_fold_change)
##            Chisq Df Pr(>Chisq)    
## Bacteria   41.45  1  1.209e-10 ***
## Wolbachia 569.67  1  < 2.2e-16 ***
## ---
## Signif. codes:  0 '***' 0.001 '**' 0.01 '*' 0.05 '.' 0.1 ' ' 1
```

```
summary(levels_systemic_DCV_model_reduced)
```

```
## Linear mixed model fit by REML ['lmerMod']
## Formula: log10(normalized_fold_change) ~ Bacteria + Wolbachia + (1 | Replicate)
##    Data: levels_systemic_DCV
## 
## REML criterion at convergence: 566.6
## 
## Scaled residuals: 
##     Min      1Q  Median      3Q     Max 
## -3.6253 -0.6301  0.1107  0.6861  2.4873 
## 
## Random effects:
##  Groups    Name        Variance Std.Dev.
##  Replicate (Intercept) 0.05906  0.2430  
##  Residual              0.90405  0.9508  
## Number of obs: 204, groups:  Replicate, 3
## 
## Fixed effects:
##                        Estimate Std. Error t value
## (Intercept)             -0.1174     0.1802  -0.652
## BacteriaA.thailandicus  -0.8770     0.1362  -6.438
## WolbachiaWolb+          -3.2667     0.1369 -23.868
## 
## Correlation of Fixed Effects:
##             (Intr) BctrA.
## BctrA.thlnd -0.391       
## WolbachWlb+ -0.374  0.148
```

- Model diagnostics

  - The residuals have a distribution close to normal

```
resid(levels_systemic_DCV_model_reduced) %>% qqnorm
```

- Average effect of *Wolbachia* and *A. thailandicus* presence

  - Decreased loads in the presence of Wolbachia and *A. thailandicus*

```
emmeans(levels_systemic_DCV_model_reduced,
        list(wolb=trt.vs.ctrl~Wolbachia,                                               Bacteria=trt.vs.ctrl~Bacteria))
```

```
## Note: Use 'contrast(regrid(object), ...)' to obtain contrasts of back-transformed estimates
## Note: Use 'contrast(regrid(object), ...)' to obtain contrasts of back-transformed estimates
```

```
## $`wolb emmeans`
##  Wolbachia emmean    SE   df lower.CL upper.CL
##  Wolb-     -0.556 0.166 2.56    -1.14   0.0262
##  Wolb+     -3.823 0.175 3.14    -4.37  -3.2789
## 
## Results are averaged over the levels of: Bacteria 
## Degrees-of-freedom method: kenward-roger 
## Results are given on the log10 (not the response) scale. 
## Confidence level used: 0.95 
## 
## $`wolb differences from control`
##  wolb.contrast     estimate    SE  df t.ratio p.value
##  (Wolb+) - (Wolb-)    -3.27 0.138 200 -23.755 <.0001 
## 
## Results are averaged over the levels of: Bacteria 
## Note: contrasts are still on the log10 scale 
## Degrees-of-freedom method: kenward-roger 
## 
## $`Bacteria emmeans`
##  Bacteria       emmean    SE   df lower.CL upper.CL
##  GF              -1.75 0.167 2.64    -2.33    -1.18
##  A.thailandicus  -2.63 0.174 3.03    -3.18    -2.08
## 
## Results are averaged over the levels of: Wolbachia 
## Degrees-of-freedom method: kenward-roger 
## Results are given on the log10 (not the response) scale. 
## Confidence level used: 0.95 
## 
## $`Bacteria differences from control`
##  Bacteria.contrast   estimate    SE  df t.ratio p.value
##  A.thailandicus - GF   -0.877 0.137 201 -6.403  <.0001 
## 
## Results are averaged over the levels of: Wolbachia 
## Note: contrasts are still on the log10 scale 
## Degrees-of-freedom method: kenward-roger
```

###### Plot

- Overall plot of all experiments

```
levels_systemic_DCV_plot_full=ggplot(levels_systemic_DCV %>% filter(Replicate=="B"))+
  aes(x=Bacteria,y=normalized_fold_change)+
  geom_beeswarm(cex = 2,aes(color=Bacteria))+
  scale_y_log10(labels=prettyNum)+
  facet_grid(~Wolbachia)+
  scale_color_manual(values=c("darkgray","black"))+
  theme_HMI()+
  stat_summary(fun="median",geom="crossbar",width=.5)+
  labs(title="Relative DCV levels",y=NULL)
  
levels_systemic_DCV_plot_full
```

- Selected panel

```
levels_systemic_DCV_plot=levels_systemic_DCV_plot_full%+%filter(levels_systemic_DCV,Replicate=="B")
levels_systemic_DCV_plot
```

###### FHV

###### Linear mixed model

- Full model

  - No interaction between microbiota and wolbachia, but a significant increase of FHV viral loads in the presence of *A.thailandicus*

```
levels_systemic_FHV_model_full=lmer(log10(normalized_fold_change)~
                                 Bacteria*Wolbachia+(1|Replicate),
                               data=levels_systemic_FHV)

Anova(levels_systemic_FHV_model_full)
```

```
## Analysis of Deviance Table (Type II Wald chisquare tests)
## 
## Response: log10(normalized_fold_change)
##                      Chisq Df Pr(>Chisq)    
## Bacteria            4.7354  1    0.02955 *  
## Wolbachia          96.5398  1    < 2e-16 ***
## Bacteria:Wolbachia  2.4587  1    0.11687    
## ---
## Signif. codes:  0 '***' 0.001 '**' 0.01 '*' 0.05 '.' 0.1 ' ' 1
```

```
summary(levels_systemic_FHV_model_full)
```

```
## Linear mixed model fit by REML ['lmerMod']
## Formula: log10(normalized_fold_change) ~ Bacteria * Wolbachia + (1 | Replicate)
##    Data: levels_systemic_FHV
## 
## REML criterion at convergence: 318.3
## 
## Scaled residuals: 
##     Min      1Q  Median      3Q     Max 
## -3.5447 -0.4986  0.0696  0.6786  2.1427 
## 
## Random effects:
##  Groups    Name        Variance Std.Dev.
##  Replicate (Intercept) 0.1286   0.3586  
##  Residual              0.5190   0.7205  
## Number of obs: 142, groups:  Replicate, 2
## 
## Fixed effects:
##                                       Estimate Std. Error t value
## (Intercept)                           -0.01433    0.27922  -0.051
## BacteriaA.thailandicus                 0.09145    0.16320   0.560
## WolbachiaWolb+                        -1.38732    0.17166  -8.082
## BacteriaA.thailandicus:WolbachiaWolb+  0.38122    0.24312   1.568
## 
## Correlation of Fixed Effects:
##             (Intr) BctrA. WlbcW+
## BctrA.thlnd -0.300              
## WolbachWlb+ -0.285  0.488       
## BctrA.t:WW+  0.201 -0.671 -0.704
```

```
levels_systemic_FHV_model_reduced=lmer(log10(normalized_fold_change)~
                                 Bacteria+Wolbachia+(1|Replicate),
                               data=levels_systemic_FHV)

Anova(levels_systemic_FHV_model_reduced)
```

```
## Analysis of Deviance Table (Type II Wald chisquare tests)
## 
## Response: log10(normalized_fold_change)
##             Chisq Df Pr(>Chisq)    
## Bacteria   4.6859  1    0.03041 *  
## Wolbachia 95.5284  1    < 2e-16 ***
## ---
## Signif. codes:  0 '***' 0.001 '**' 0.01 '*' 0.05 '.' 0.1 ' ' 1
```

```
summary(levels_systemic_FHV_model_reduced)
```

```
## Linear mixed model fit by REML ['lmerMod']
## Formula: log10(normalized_fold_change) ~ Bacteria + Wolbachia + (1 | Replicate)
##    Data: levels_systemic_FHV
## 
## REML criterion at convergence: 319.8
## 
## Scaled residuals: 
##     Min      1Q  Median      3Q     Max 
## -3.6661 -0.4615  0.1187  0.6816  2.2533 
## 
## Random effects:
##  Groups    Name        Variance Std.Dev.
##  Replicate (Intercept) 0.1288   0.3589  
##  Residual              0.5245   0.7242  
## Number of obs: 142, groups:  Replicate, 2
## 
## Fixed effects:
##                        Estimate Std. Error t value
## (Intercept)             -0.1024     0.2739  -0.374
## BacteriaA.thailandicus   0.2632     0.1216   2.165
## WolbachiaWolb+          -1.1978     0.1226  -9.774
## 
## Correlation of Fixed Effects:
##             (Intr) BctrA.
## BctrA.thlnd -0.228       
## WolbachWlb+ -0.207  0.028
```

- Close to linear distribution of residuals

```
resid(levels_systemic_FHV_model_reduced) %>% qqnorm
```

- Average effect of *Wolbachia* and *A. thailandicus* presence

  - Decreased loads in the presence of Wolbachia
  - Increased loads in the presence of *A. thailandicus*

```
emmeans(levels_systemic_FHV_model_reduced,list(
  Bacteria=pairwise~Bacteria,
  wolb=pairwise~Wolbachia
))
```

```
## Note: Use 'contrast(regrid(object), ...)' to obtain contrasts of back-transformed estimates
## Note: Use 'contrast(regrid(object), ...)' to obtain contrasts of back-transformed estimates
```

```
## $`Bacteria emmeans`
##  Bacteria       emmean    SE   df lower.CL upper.CL
##  GF             -0.701 0.268 1.11    -3.40     2.00
##  A.thailandicus -0.438 0.268 1.11    -3.13     2.25
## 
## Results are averaged over the levels of: Wolbachia 
## Degrees-of-freedom method: kenward-roger 
## Results are given on the log10 (not the response) scale. 
## Confidence level used: 0.95 
## 
## $`Bacteria pairwise differences`
##  Bacteria.contrast   estimate    SE  df t.ratio p.value
##  GF - A.thailandicus   -0.263 0.122 138 -2.165  0.0321 
## 
## Results are averaged over the levels of: Wolbachia 
## Note: contrasts are still on the log10 scale 
## Degrees-of-freedom method: kenward-roger 
## 
## $`wolb emmeans`
##  Wolbachia  emmean    SE   df lower.CL upper.CL
##  Wolb-      0.0292 0.267 1.09    -2.77     2.82
##  Wolb+     -1.1686 0.270 1.14    -3.75     1.42
## 
## Results are averaged over the levels of: Bacteria 
## Degrees-of-freedom method: kenward-roger 
## Results are given on the log10 (not the response) scale. 
## Confidence level used: 0.95 
## 
## $`wolb pairwise differences`
##  wolb.contrast     estimate    SE  df t.ratio p.value
##  (Wolb-) - (Wolb+)      1.2 0.123 138 9.767   <.0001 
## 
## Results are averaged over the levels of: Bacteria 
## Note: contrasts are still on the log10 scale 
## Degrees-of-freedom method: kenward-roger
```

###### Plot

```
levels_systemic_FHV_plot_full=ggplot(levels_systemic_FHV %>% filter(Replicate=="A"))+
  aes(x=Bacteria,y=normalized_fold_change)+
  geom_beeswarm(cex = 2,aes(color=Bacteria))+
  scale_y_log10(labels=prettyNum)+
  facet_grid(~Wolbachia)+
  scale_color_manual(values=c("darkgray","black"))+
  theme_HMI()+
  stat_summary(fun="median",geom="crossbar",width=.5)+
  labs(title="Relative FHV levels",y=NULL)

levels_systemic_FHV_plot_full
```

- Selected panel

```
levels_systemic_FHV_plot=levels_systemic_FHV_plot_full%+%filter(levels_systemic_FHV,Replicate=="A")
levels_systemic_FHV_plot
```

##### Arranged plot

```
fig1_plotlist=list(surv_systemic_DCV_panel
  ,surv_systemic_DCV_hazards_plot+facet_grid(~"") # small hack to align the x-axis
  ,levels_systemic_DCV_plot+guides(colour="none")
  ,surv_systemic_FHV_panel 
  , surv_systemic_FHV_hazards_plot+facet_grid(~"") # small hack to align the x-axis 
  ,levels_systemic_FHV_plot+guides(color="none"))

fig1=wrap_plots(fig1_plotlist,guides = "collect")+
  plot_annotation(tag_levels = "A")&
  theme(legend.position = "bottom",plot.tag = element_text(face="bold"))

fig1
```

```
cowplot::ggsave2(here("manuscript","fig1.png"),
                 plot = fig1,
                 width = 18,height = 9,units = "cm",scale=1.5)
```

#### Figure S1 - DCV in Oregon R

##### Survival

###### DCV

###### Data loading

```
surv_systemic_OR_DCV=filter(surv_systemic,Virus%in%c("DCV"),
                            Stock=="Oregon R")
```

###### Mixed effects Cox model

- Significant effect of having Bacteria microbiota, Wolbachia, Sex and Experiment
- *A.thai* significantly increases the risk of death, with or without wolbachia

```
surv_systemic_OR_DCV_model_mixed_full=coxme(Surv(day_status,status)~
                                              Wolbachia*Bacteria*Sex+
                                              (1|Experiment)+
                                              (1|RepFull),
                                         surv_systemic_OR_DCV)
Anova(surv_systemic_OR_DCV_model_mixed_full)
```

```
## Analysis of Deviance Table (Type II tests)
## 
## Response: Surv(day_status, status)
##                        Df    Chisq Pr(>Chisq)    
## Wolbachia               1 304.3087  < 2.2e-16 ***
## Bacteria                1  17.9928  2.217e-05 ***
## Sex                     1   7.2589   0.007055 ** 
## Wolbachia:Bacteria      1   1.4223   0.233020    
## Wolbachia:Sex           1   5.9261   0.014918 *  
## Bacteria:Sex            1   0.0543   0.815679    
## Wolbachia:Bacteria:Sex  1   0.0990   0.752989    
## ---
## Signif. codes:  0 '***' 0.001 '**' 0.01 '*' 0.05 '.' 0.1 ' ' 1
```

```
summary(surv_systemic_OR_DCV_model_mixed_full)
```

```
## Cox mixed-effects model fit by maximum likelihood
##   Data: surv_systemic_OR_DCV
##   events, n = 406, 600
##   Iterations= 10 64 
##                     NULL Integrated   Fitted
## Log-likelihood -2410.758  -2162.113 -2116.29
## 
##                    Chisq    df p    AIC    BIC
## Integrated loglik 497.29  9.00 0 479.29 443.23
##  Penalized loglik 588.93 44.23 0 500.48 323.28
## 
## Model:  Surv(day_status, status) ~ Wolbachia * Bacteria * Sex + (1 |      Experiment) + (1 | RepFull) 
## Fixed coefficients
##                                                       coef  exp(coef)  se(coef)
## WolbachiaWolb+                                 -3.37605368 0.03418208 0.3397848
## BacteriaA.thailandicus                          0.37879613 1.46052524 0.2211565
## SexMales                                        0.06183625 1.06378813 0.2229254
## WolbachiaWolb+:BacteriaA.thailandicus           0.43075907 1.53842485 0.4236294
## WolbachiaWolb+:SexMales                         0.77507075 2.17074570 0.4248128
## BacteriaA.thailandicus:SexMales                 0.11651916 1.12357904 0.3148241
## WolbachiaWolb+:BacteriaA.thailandicus:SexMales -0.17536309 0.83915227 0.5572376
##                                                    z     p
## WolbachiaWolb+                                 -9.94 0.000
## BacteriaA.thailandicus                          1.71 0.087
## SexMales                                        0.28 0.780
## WolbachiaWolb+:BacteriaA.thailandicus           1.02 0.310
## WolbachiaWolb+:SexMales                         1.82 0.068
## BacteriaA.thailandicus:SexMales                 0.37 0.710
## WolbachiaWolb+:BacteriaA.thailandicus:SexMales -0.31 0.750
## 
## Random effects
##  Group      Variable  Std Dev    Variance  
##  Experiment Intercept 0.22574821 0.05096225
##  RepFull    Intercept 0.39830320 0.15864544
```

- Reduced model

```
surv_systemic_OR_DCV_model_mixed_reduced=coxme(Surv(day_status,status)~
                                                 Wolbachia+Bacteria+Sex+
                                                 (1|Experiment)+(1|RepFull),
                                         surv_systemic_OR_DCV)
Anova(surv_systemic_OR_DCV_model_mixed_reduced)
```

```
## Analysis of Deviance Table (Type II tests)
## 
## Response: Surv(day_status, status)
##           Df    Chisq Pr(>Chisq)    
## Wolbachia  1 303.8373  < 2.2e-16 ***
## Bacteria   1  17.4016  3.026e-05 ***
## Sex        1   7.5056   0.006151 ** 
## ---
## Signif. codes:  0 '***' 0.001 '**' 0.01 '*' 0.05 '.' 0.1 ' ' 1
```

- Average effect of *Wolbachia* and *A. thailandicus* presence
- Decreased risk in the presence of Wolbachia and increased risk in the presence of *A. thailandicus*

```
surv_systemic_OR_DCV_emmeans=emmeans(surv_systemic_OR_DCV_model_mixed_reduced,                                       list(Bacteria=~Bacteria,
                                                                                                                          wolb=~Wolbachia))

surv_systemic_OR_DCV_emmeans %>% 
  contrast("trt.vs.ctrl")
```

```
## $Bacteria
##  contrast            estimate    SE  df z.ratio p.value
##  A.thailandicus - GF    0.556 0.133 Inf 4.172   <.0001 
## 
## Results are averaged over the levels of: Wolbachia, Sex 
## Results are given on the log (not the response) scale. 
## 
## $wolb
##  contrast          estimate    SE  df z.ratio p.value
##  (Wolb+) - (Wolb-)    -2.78 0.159 Inf -17.431 <.0001 
## 
## Results are averaged over the levels of: Bacteria, Sex 
## Results are given on the log (not the response) scale.
```

###### Plots

- Overall plot of all experiments

```
surv_systemic_OR_DCV_fit=survfit(Surv(day_status,status)~Bacteria+Wolbachia+Experiment+Sex,surv_systemic_OR_DCV)

surv_systemic_OR_DCV_summary=surv_summary(surv_systemic_OR_DCV_fit,data=surv_systemic_OR_DCV) %>% 
  mutate(Bacteria=fct_relevel(Bacteria,"GF")) %>%
  group_by(strata,Bacteria,Sex,Wolbachia,Experiment) %>% 
  group_modify(~{
    if(min(.x$time)!=0){.x=add_row(.x,time=0,surv=1)}else{.x}})

surv_systemic_OR_DCV_plot=ggplot(surv_systemic_OR_DCV_summary)+
  aes(x=time,y=surv)+
  geom_line(aes(linetype=Wolbachia,color=Bacteria))+
  geom_point(aes(color=Bacteria))+
  scale_color_manual(values=c("darkgray","black"))+
  scale_y_continuous(limits=c(0,1))+
  facet_grid(Sex~Experiment)+
  labs(title="Proportion of surviving individuals after DCV infection",y=NULL)+
  theme_HMI()
  
surv_systemic_OR_DCV_plot
```

- Hazard ratio plot

```
surv_systemic_OR_DCV_hazards=map_dfr(surv_systemic_OR_DCV_emmeans,{~
                                   contrast(.,"trt.vs.ctrl") %>% 
                                     confint%>% data.frame}) 


surv_systemic_OR_DCV_hazards_plot=ggplot(surv_systemic_OR_DCV_hazards)+
  aes(x=contrast,y=estimate)+
  geom_bar(aes(x=contrast,y=estimate),stat="identity")+
  geom_errorbar(aes(ymin=asymp.LCL,ymax=asymp.UCL),width=.5)+
  theme_HMI() +
  labs(title="Cox regression coefficients",y=NULL,x="Contrast")


surv_systemic_OR_DCV_hazards_plot
```

###### FHV

###### Data loading

```
surv_systemic_OR_FHV=filter(surv_systemic,Virus=="FHV",Stock=="Oregon R")
```

###### Mixed effects Cox model

- Full model

  - As for DCV, effect of having A. thailandicus\*. Interaction with wolbachia and Sex of the flies
  - *A.thai* increases the risk of death, especially in the presence of Wolbachia.

```
surv_systemic_OR_FHV_model_mixed_full=coxme(Surv(day_status,status)~
                                              Wolbachia*Bacteria*Sex+
                                              (1|Experiment)+(1|RepFull), surv_systemic_OR_FHV)
Anova(surv_systemic_OR_FHV_model_mixed_full)
```

```
## Analysis of Deviance Table (Type II tests)
## 
## Response: Surv(day_status, status)
##                        Df    Chisq Pr(>Chisq)    
## Wolbachia               1 130.5272  < 2.2e-16 ***
## Bacteria                1  19.2542  1.144e-05 ***
## Sex                     1  12.0226  0.0005256 ***
## Wolbachia:Bacteria      1   8.2999  0.0039648 ** 
## Wolbachia:Sex           1   0.3034  0.5817392    
## Bacteria:Sex            1  10.4815  0.0012057 ** 
## Wolbachia:Bacteria:Sex  1   0.4265  0.5137088    
## ---
## Signif. codes:  0 '***' 0.001 '**' 0.01 '*' 0.05 '.' 0.1 ' ' 1
```

```
summary(surv_systemic_OR_FHV_model_mixed_full)
```

```
## Cox mixed-effects model fit by maximum likelihood
##   Data: surv_systemic_OR_FHV
##   events, n = 371, 600
##   Iterations= 12 75 
##                     NULL Integrated    Fitted
## Log-likelihood -2223.317  -2107.102 -2079.869
## 
##                    Chisq    df p    AIC    BIC
## Integrated loglik 232.43  9.00 0 214.43 179.18
##  Penalized loglik 286.89 29.58 0 227.73 111.87
## 
## Model:  Surv(day_status, status) ~ Wolbachia * Bacteria * Sex + (1 |      Experiment) + (1 | RepFull) 
## Fixed coefficients
##                                                      coef exp(coef)  se(coef)
## WolbachiaWolb+                                 -2.0518853 0.1284924 0.2595794
## BacteriaA.thailandicus                         -0.1264890 0.8811839 0.2033429
## SexMales                                       -0.9785505 0.3758555 0.2163417
## WolbachiaWolb+:BacteriaA.thailandicus           0.8831076 2.4184035 0.3412784
## WolbachiaWolb+:SexMales                         0.3460943 1.4135358 0.4053828
## BacteriaA.thailandicus:SexMales                 0.9047023 2.4711961 0.3005473
## WolbachiaWolb+:BacteriaA.thailandicus:SexMales -0.3362418 0.7144503 0.5148605
##                                                    z       p
## WolbachiaWolb+                                 -7.90 2.7e-15
## BacteriaA.thailandicus                         -0.62 5.3e-01
## SexMales                                       -4.52 6.1e-06
## WolbachiaWolb+:BacteriaA.thailandicus           2.59 9.7e-03
## WolbachiaWolb+:SexMales                         0.85 3.9e-01
## BacteriaA.thailandicus:SexMales                 3.01 2.6e-03
## WolbachiaWolb+:BacteriaA.thailandicus:SexMales -0.65 5.1e-01
## 
## Random effects
##  Group      Variable  Std Dev    Variance  
##  Experiment Intercept 0.34205103 0.11699891
##  RepFull    Intercept 0.28161973 0.07930967
```

- Reduced model

```
surv_systemic_OR_FHV_model_mixed_reduced=coxme(Surv(day_status,status)~
                                                 Bacteria*(Wolbachia+Sex)+
                                                 (1|Experiment)+(1|RepFull),
                                         surv_systemic_OR_FHV)
Anova(surv_systemic_OR_FHV_model_mixed_reduced)
```

```
## Analysis of Deviance Table (Type II tests)
## 
## Response: Surv(day_status, status)
##                    Df   Chisq Pr(>Chisq)    
## Bacteria            1  19.393  1.064e-05 ***
## Wolbachia           1 129.778  < 2.2e-16 ***
## Sex                 1  11.880  0.0005674 ***
## Bacteria:Wolbachia  1   8.760  0.0030790 ** 
## Bacteria:Sex        1  11.272  0.0007870 ***
## ---
## Signif. codes:  0 '***' 0.001 '**' 0.01 '*' 0.05 '.' 0.1 ' ' 1
```

```
summary(surv_systemic_OR_FHV_model_mixed_reduced)
```

```
## Cox mixed-effects model fit by maximum likelihood
##   Data: surv_systemic_OR_FHV
##   events, n = 371, 600
##   Iterations= 14 73 
##                     NULL Integrated    Fitted
## Log-likelihood -2223.317  -2107.463 -2080.162
## 
##                    Chisq    df p    AIC    BIC
## Integrated loglik 231.71  7.00 0 217.71 190.30
##  Penalized loglik 286.31 27.99 0 230.32 120.69
## 
## Model:  Surv(day_status, status) ~ Bacteria * (Wolbachia + Sex) + (1 |      Experiment) + (1 | RepFull) 
## Fixed coefficients
##                                              coef exp(coef)  se(coef)     z
## BacteriaA.thailandicus                -0.08595009 0.9176400 0.1867383 -0.46
## WolbachiaWolb+                        -1.91737414 0.1469924 0.2023653 -9.47
## SexMales                              -0.88253436 0.4137330 0.1842264 -4.79
## BacteriaA.thailandicus:WolbachiaWolb+  0.75449428 2.1265358 0.2549193  2.96
## BacteriaA.thailandicus:SexMales        0.81290991 2.2544587 0.2421305  3.36
##                                             p
## BacteriaA.thailandicus                6.5e-01
## WolbachiaWolb+                        0.0e+00
## SexMales                              1.7e-06
## BacteriaA.thailandicus:WolbachiaWolb+ 3.1e-03
## BacteriaA.thailandicus:SexMales       7.9e-04
## 
## Random effects
##  Group      Variable  Std Dev    Variance  
##  Experiment Intercept 0.34146214 0.11659639
##  RepFull    Intercept 0.28214640 0.07960659
```

- Average effect of *Wolbachia* and *A. thailandicus* presence
- Decreased risk in the presence of Wolbachia and increased risk in the presence of *A. thailandicus*

```
surv_systemic_OR_FHV_emmeans=emmeans(surv_systemic_OR_FHV_model_mixed_reduced,                                       list(Bacteria=~Bacteria,
                                                                                                                          wolb=~Wolbachia))
```

```
## NOTE: Results may be misleading due to involvement in interactions
## NOTE: Results may be misleading due to involvement in interactions
```

```
surv_systemic_OR_FHV_emmeans %>% 
  contrast("trt.vs.ctrl")
```

```
## $Bacteria
##  contrast            estimate    SE  df z.ratio p.value
##  A.thailandicus - GF    0.698 0.129 Inf 5.422   <.0001 
## 
## Results are averaged over the levels of: Wolbachia, Sex 
## Results are given on the log (not the response) scale. 
## 
## $wolb
##  contrast          estimate    SE  df z.ratio p.value
##  (Wolb+) - (Wolb-)    -1.54 0.131 Inf -11.766 <.0001 
## 
## Results are averaged over the levels of: Bacteria, Sex 
## Results are given on the log (not the response) scale.
```

```
surv_systemic_OR_FHV_emmeans_full=emmeans(surv_systemic_OR_FHV_model_mixed_reduced,                                       list(Bacteria=~Bacteria|Wolbachia,
                                                                                                                          wolb=~Wolbachia|Bacteria))
 surv_systemic_OR_FHV_emmeans_full %>% 
    contrast("trt.vs.ctrl")
```

```
## $Bacteria
## Wolbachia = Wolb-:
##  contrast            estimate    SE  df z.ratio p.value
##  A.thailandicus - GF    0.321 0.149 Inf 2.149   0.0316 
## 
## Wolbachia = Wolb+:
##  contrast            estimate    SE  df z.ratio p.value
##  A.thailandicus - GF    1.075 0.208 Inf 5.162   <.0001 
## 
## Results are averaged over the levels of: Sex 
## Results are given on the log (not the response) scale. 
## 
## $wolb
## Bacteria = GF:
##  contrast          estimate    SE  df z.ratio p.value
##  (Wolb+) - (Wolb-)    -1.92 0.202 Inf -9.475  <.0001 
## 
## Bacteria = A.thailandicus:
##  contrast          estimate    SE  df z.ratio p.value
##  (Wolb+) - (Wolb-)    -1.16 0.161 Inf -7.239  <.0001 
## 
## Results are averaged over the levels of: Sex 
## Results are given on the log (not the response) scale.
```

###### Plots

- Overall plot of all experiments

```
surv_systemic_OR_FHV_fit=survfit(Surv(day_status,status)~Bacteria+Wolbachia+Experiment+Sex,surv_systemic_OR_FHV)

surv_systemic_OR_FHV_summary=surv_summary(surv_systemic_OR_FHV_fit,data=surv_systemic_OR_FHV) %>% 
  mutate(Bacteria=fct_relevel(Bacteria,"GF")) %>% 
  group_by(strata,Bacteria,Wolbachia,Sex,Experiment) %>% 
  group_modify(~{
    if(min(.x$time)!=0){.x=add_row(.x,time=0,surv=1)}else{.x}})

surv_systemic_OR_FHV_plot=ggplot(surv_systemic_OR_FHV_summary)+
  aes(x=time,y=surv)+
  geom_line(aes(linetype=Wolbachia,color=Bacteria))+
  geom_point(aes(color=Bacteria))+
  scale_color_manual(values=c("darkgray","black"))+
  scale_y_continuous(limits=c(0,1))+
  facet_grid(Sex~Experiment)+
  labs(title="Proportion of surviving individuals after FHV infection",y=NULL)+
  theme_HMI()
  
surv_systemic_OR_FHV_plot
```

- Hazard ratio

```
surv_systemic_OR_FHV_hazards=map_dfr(surv_systemic_OR_FHV_emmeans,{~
                                   contrast(.,"trt.vs.ctrl") %>% 
                                     confint%>% data.frame}) 


surv_systemic_OR_FHV_hazards_plot=ggplot(surv_systemic_OR_FHV_hazards)+
  aes(x=contrast,y=estimate)+
  geom_bar(aes(x=contrast,y=estimate),stat="identity")+
  geom_errorbar(aes(ymin=asymp.LCL,ymax=asymp.UCL),width=.5)+
  theme_HMI() +
  labs(title="Cox regression coefficients",y=NULL,x="Contrast")


surv_systemic_OR_FHV_hazards_plot
```

##### Arranged plot

```
or_plotlist=list(surv_systemic_OR_DCV_plot%+%
  filter(surv_systemic_OR_DCV_summary,Experiment=="C")+
    facet_grid(~Sex)
  ,surv_systemic_OR_DCV_hazards_plot+facet_grid(~"")
  ,surv_systemic_OR_FHV_plot%+%
  filter(surv_systemic_OR_FHV_summary,Experiment=="C")+
    facet_grid(~Sex)
  ,surv_systemic_OR_FHV_hazards_plot+facet_grid(~""))

fig_s1=wrap_plots(or_plotlist,guides="collect")+ 
  plot_annotation(tag_levels = "A")&
  theme(legend.position = "bottom",plot.tag=element_text(face="bold"))

fig_s1
```

```
ggsave2(plot=fig_s1,here("manuscript","figS1.png"),
        width = 14,height=9,units="cm",scale=1.5)
```

#### Figure 2 - DCV with other bacteria

##### Survival

###### Data loading

```
surv_systemic_multi_DCV=surv_systemic_multi %>% 
  filter(Virus=="DCV") %>% 
  mutate(L_brevis=fct_collapse(Bacteria,"Yes"=str_subset(Bacteria,"brevis"),other_level = "No")
         ,A_thai=fct_collapse(Bacteria,"Yes"=str_subset(Bacteria,"thai"),other_level = "No")
         )
```

###### Mixed effect Cox Model

- The effect on survival of colonization by *A. thai* or *L. brevis* are tested independently.
- There is a significant deleterious effect of *A. thai* in survival, whereas *L. brevis* has no effect.

```
surv_systemic_multi_DCV_model_mixed_bybac_full=coxme(Surv(day_status,status)~
                                                  L_brevis*A_thai+
                                                    (1|Experiment)+
                                                    (1|RepFull),
                                                
                                         surv_systemic_multi_DCV)
Anova(surv_systemic_multi_DCV_model_mixed_bybac_full)
```

```
## Analysis of Deviance Table (Type II tests)
## 
## Response: Surv(day_status, status)
##                 Df  Chisq Pr(>Chisq)   
## L_brevis         1 0.1912    0.66191   
## A_thai           1 9.7224    0.00182 **
## L_brevis:A_thai  1 2.0214    0.15510   
## ---
## Signif. codes:  0 '***' 0.001 '**' 0.01 '*' 0.05 '.' 0.1 ' ' 1
```

```
summary(surv_systemic_multi_DCV_model_mixed_bybac_full)
```

```
## Cox mixed-effects model fit by maximum likelihood
##   Data: surv_systemic_multi_DCV
##   events, n = 288, 300
##   Iterations= 13 68 
##                     NULL Integrated    Fitted
## Log-likelihood -1394.919  -1383.191 -1362.276
## 
##                   Chisq    df          p   AIC    BIC
## Integrated loglik 23.46  5.00 2.7610e-04 13.46  -4.86
##  Penalized loglik 65.29 20.39 1.3445e-06 24.51 -50.18
## 
## Model:  Surv(day_status, status) ~ L_brevis * A_thai + (1 | Experiment) +      (1 | RepFull) 
## Fixed coefficients
##                           coef exp(coef)  se(coef)     z    p
## L_brevisNo           0.1378628 1.1478181 0.2010706  0.69 0.49
## A_thaiNo            -0.2464636 0.7815598 0.2029333 -1.21 0.22
## L_brevisNo:A_thaiNo -0.4066359 0.6658866 0.2860111 -1.42 0.16
## 
## Random effects
##  Group      Variable  Std Dev    Variance  
##  Experiment Intercept 0.20006206 0.04002483
##  RepFull    Intercept 0.29998386 0.08999032
```

- Reduced model

```
surv_systemic_multi_DCV_model_mixed_bybac_reduced=coxme(Surv(day_status,status)~
                                                  L_brevis+A_thai+
                                                    (1|Experiment)+(1|RepFull),
                                                
                                         surv_systemic_multi_DCV)
Anova(surv_systemic_multi_DCV_model_mixed_bybac_reduced)
```

```
## Analysis of Deviance Table (Type II tests)
## 
## Response: Surv(day_status, status)
##          Df  Chisq Pr(>Chisq)   
## L_brevis  1 0.1849   0.667169   
## A_thai    1 9.5005   0.002054 **
## ---
## Signif. codes:  0 '***' 0.001 '**' 0.01 '*' 0.05 '.' 0.1 ' ' 1
```

```
summary(surv_systemic_multi_DCV_model_mixed_bybac_reduced)
```

```
## Cox mixed-effects model fit by maximum likelihood
##   Data: surv_systemic_multi_DCV
##   events, n = 288, 300
##   Iterations= 16 67 
##                     NULL Integrated    Fitted
## Log-likelihood -1394.919  -1384.185 -1361.626
## 
##                   Chisq    df          p   AIC    BIC
## Integrated loglik 21.47  4.00 2.5571e-04 13.47  -1.18
##  Penalized loglik 66.58 20.98 1.2097e-06 24.63 -52.21
## 
## Model:  Surv(day_status, status) ~ L_brevis + A_thai + (1 | Experiment) +      (1 | RepFull) 
## Fixed coefficients
##                   coef exp(coef)  se(coef)     z      p
## L_brevisNo -0.06274695 0.9391811 0.1459106 -0.43 0.6700
## A_thaiNo   -0.45236934 0.6361192 0.1467639 -3.08 0.0021
## 
## Random effects
##  Group      Variable  Std Dev    Variance  
##  Experiment Intercept 0.19869151 0.03947832
##  RepFull    Intercept 0.31705106 0.10052137
```

- Average effect of *A. thailandicus* and *L. brevis* presence
- Increased risk in the presence of *A. thailandicus*

```
surv_systemic_multi_DCV_emmeans=emmeans(surv_systemic_multi_DCV_model_mixed_bybac_reduced,
        list(thai=~A_thai,lact=~L_brevis))

surv_systemic_multi_DCV_emmeans %>% 
  contrast("trt.vs.ctrl")
```

```
## $thai
##  contrast estimate    SE  df z.ratio p.value
##  No - Yes   -0.452 0.147 Inf -3.082  0.0021 
## 
## Results are averaged over the levels of: L_brevis 
## Results are given on the log (not the response) scale. 
## 
## $lact
##  contrast estimate    SE  df z.ratio p.value
##  No - Yes  -0.0627 0.146 Inf -0.430  0.6672 
## 
## Results are averaged over the levels of: A_thai 
## Results are given on the log (not the response) scale.
```

###### Plots

- Overall plot of all experiments

```
surv_systemic_multi_DCV_fit=survfit(Surv(day_status,status)~Bacteria+Wolbachia+Experiment+Sex,surv_systemic_multi_DCV)

surv_systemic_multi_DCV_summary=surv_summary(surv_systemic_multi_DCV_fit,data=surv_systemic_multi_DCV) %>% 
  mutate(Bacteria=fct_relevel(Bacteria,"GF")) %>% 
  group_by(strata,Bacteria,Wolbachia,Sex,Experiment) %>% 
  group_modify(~{
    if(min(.x$time)!=0){.x=add_row(.x,time=0,surv=1)}else{.x}})

surv_systemic_multi_DCV_plot=ggplot(surv_systemic_multi_DCV_summary)+
  aes(x=time,y=surv)+
  geom_line(aes(color=Bacteria))+
  geom_point(aes(color=Bacteria,shape=Bacteria))+
  scale_color_manual(values=c("darkgray","black","black","black"))+
  scale_y_continuous(limits=c(0,1))+
  facet_grid(Sex~Experiment)+
  labs(title="Proportion of surviving individuals after DCV infection",x="Time (d)",y=NULL)+
  theme_HMI()
  
surv_systemic_multi_DCV_plot
```

- Selected panel

```
surv_systemic_multi_DCV_panel=
  surv_systemic_multi_DCV_plot %+% 
  filter(surv_systemic_multi_DCV_summary,Experiment=="A")+
  facet_null()
```

- Hazard ratio plot

```
surv_systemic_multi_DCV_emm_bybac=emmeans(surv_systemic_multi_DCV_model_mixed_bybac_reduced,
                                          list("L.brevis (- vs +)"=~L_brevis,
                                                                                                 "A.thailandicus (- vs +)"=~A_thai)) %>% 
  contrast("pairwise") %>% 
  map_dfr(~{confint(.) %>% data.frame},.id = "Bacteria")  
  

surv_systemic_multi_DCV_hazard_plot=ggplot(surv_systemic_multi_DCV_emm_bybac) +
  aes(x=Bacteria,y=estimate,ymin=asymp.LCL,ymax=asymp.UCL)+
  geom_bar(stat="identity")+
  geom_errorbar(width=.5)+
  theme_HMI()+
  labs(title="Cox regression coefficients",y=NULL,x="Contrast")
  
surv_systemic_multi_DCV_hazard_plot
```

##### Arranged plot

```
fig2=surv_systemic_multi_DCV_panel+surv_systemic_multi_DCV_hazard_plot+
  patchwork::plot_annotation(tag_levels = "A")&
  theme(legend.position = "bottom",plot.tag = element_text(face="bold"))

fig2
```

```
ggsave2(filename = here("manuscript","fig2.png"),plot = fig2,
        width=12,
        height=6,
        units="cm",scale=2)
```

#### Figure 3 - Oral Infection

##### Survival

###### Data loading

```
surv_oral_DCV=filter(surv_oral,Virus%in%c("DCV")) %>% 
  mutate(
  L_brevis=fct_collapse(Bacteria,"Yes"=str_subset(Bacteria,"brevis"),other_level = "No")
         ,A_thai=fct_collapse(Bacteria,"Yes"=str_subset(Bacteria,"thai"),other_level = "No")
         ) %>% 
  filter(Virus=="DCV")
```

###### Mixed effect Cox model

- Full model
- No effect of the any bacteria

```
surv_oral_DCV_model_mixed_full=coxme(Surv(day_status,status)~
                                       L_brevis*A_thai*Sex+
                                       (1|Experiment)+(1|RepFull),
                                         surv_oral_DCV)


Anova(surv_oral_DCV_model_mixed_full)
```

```
## Analysis of Deviance Table (Type II tests)
## 
## Response: Surv(day_status, status)
##                     Df   Chisq Pr(>Chisq)    
## L_brevis             1  1.1114    0.29179    
## A_thai               1  0.0558    0.81321    
## Sex                  1 21.5290  3.485e-06 ***
## L_brevis:A_thai      1  0.4145    0.51970    
## L_brevis:Sex         1  2.9473    0.08602 .  
## A_thai:Sex           1  0.5606    0.45400    
## L_brevis:A_thai:Sex  1  0.4230    0.51544    
## ---
## Signif. codes:  0 '***' 0.001 '**' 0.01 '*' 0.05 '.' 0.1 ' ' 1
```

```
summary(surv_oral_DCV_model_mixed_full)
```

```
## Cox mixed-effects model fit by maximum likelihood
##   Data: surv_oral_DCV
##   events, n = 713, 1155
##   Iterations= 12 64 
##                     NULL Integrated    Fitted
## Log-likelihood -4740.014  -4702.217 -4631.594
## 
##                    Chisq    df          p   AIC     BIC
## Integrated loglik  75.59  9.00 1.2059e-12 57.59   16.47
##  Penalized loglik 216.84 62.58 0.0000e+00 91.67 -194.30
## 
## Model:  Surv(day_status, status) ~ L_brevis * A_thai * Sex + (1 | Experiment) +      (1 | RepFull) 
## Fixed coefficients
##                                      coef exp(coef)  se(coef)     z       p
## L_brevisNo                   -0.280168780 0.7556562 0.2045739 -1.37 0.17000
## A_thaiNo                     -0.095925760 0.9085315 0.2046827 -0.47 0.64000
## SexMales                     -0.828119057 0.4368702 0.2153589 -3.85 0.00012
## L_brevisNo:A_thaiNo          -0.005955477 0.9940622 0.2922011 -0.02 0.98000
## L_brevisNo:SexMales           0.503740658 1.6549001 0.3011800  1.67 0.09400
## A_thaiNo:SexMales             0.298049244 1.3472281 0.3013116  0.99 0.32000
## L_brevisNo:A_thaiNo:SexMales -0.276449562 0.7584719 0.4250523 -0.65 0.52000
## 
## Random effects
##  Group      Variable  Std Dev     Variance   
##  Experiment Intercept 0.079594312 0.006335255
##  RepFull    Intercept 0.407635106 0.166166380
```

- Reduced model

```
surv_oral_DCV_model_mixed_reduced=coxme(Surv(day_status,status)~
                                       L_brevis+A_thai+Sex+
                                       (1|Experiment)+(1|RepFull),
                                         surv_oral_DCV)
Anova(surv_oral_DCV_model_mixed_reduced)
```

```
## Analysis of Deviance Table (Type II tests)
## 
## Response: Surv(day_status, status)
##          Df   Chisq Pr(>Chisq)    
## L_brevis  1  1.0678     0.3014    
## A_thai    1  0.0498     0.8234    
## Sex       1 21.0507  4.473e-06 ***
## ---
## Signif. codes:  0 '***' 0.001 '**' 0.01 '*' 0.05 '.' 0.1 ' ' 1
```

```
summary(surv_oral_DCV_model_mixed_reduced)
```

```
## Cox mixed-effects model fit by maximum likelihood
##   Data: surv_oral_DCV
##   events, n = 713, 1155
##   Iterations= 11 48 
##                     NULL Integrated   Fitted
## Log-likelihood -4740.014   -4704.35 -4630.82
## 
##                    Chisq    df          p   AIC     BIC
## Integrated loglik  71.33  5.00 5.4179e-14 61.33   38.48
##  Penalized loglik 218.39 62.48 0.0000e+00 93.43 -192.05
## 
## Model:  Surv(day_status, status) ~ L_brevis + A_thai + Sex + (1 | Experiment) +      (1 | RepFull) 
## Fixed coefficients
##                   coef exp(coef)  se(coef)     z       p
## L_brevisNo -0.11143304 0.8945513 0.1078352 -1.03 3.0e-01
## A_thaiNo   -0.02406218 0.9762250 0.1078010 -0.22 8.2e-01
## SexMales   -0.49607275 0.6089173 0.1081216 -4.59 4.5e-06
## 
## Random effects
##  Group      Variable  Std Dev     Variance   
##  Experiment Intercept 0.075386263 0.005683089
##  RepFull    Intercept 0.421464585 0.177632396
```

###### Plots

- Overall plot of all experiments

```
surv_oral_DCV_fit=survfit(Surv(day_status,status)~Bacteria+Wolbachia+Experiment+Sex,surv_oral_DCV)

surv_oral_DCV_summary=surv_summary(surv_oral_DCV_fit,data=surv_oral_DCV) %>% 
  mutate(Bacteria=fct_relevel(Bacteria,!!!levels(surv_oral_DCV$Bacteria))) %>% 
  group_by(strata,Bacteria,Wolbachia,Sex,Experiment) %>% 
  group_modify(~{
    if(min(.x$time)!=0){.x=add_row(.x,time=0,surv=1)}else{.x}})

surv_oral_DCV_plot=ggplot(surv_oral_DCV_summary)+
  aes(x=time,y=surv)+
  geom_line(aes(color=Bacteria))+
  geom_point(aes(color=Bacteria,shape=Bacteria))+
  scale_color_manual(values=c("darkgray","black","black","black"))+
  scale_y_continuous(limits=c(0,1))+
  facet_grid(Sex~Experiment)+
  labs(title="Proportion of surviving individuals after DCV infection",x="Time (d)",y=NULL)+
  theme_HMI()
  
surv_oral_DCV_plot
```

- Selected panel

```
surv_oral_DCV_panel=
  surv_oral_DCV_plot %+% 
  filter(surv_oral_DCV_summary,Experiment=="C")+
  facet_grid(~Sex)
  
surv_oral_DCV_panel
```

- Hazard Ratio plot

```
surv_oral_DCV_emm_bybac=emmeans(surv_oral_DCV_model_mixed_reduced,
                                list("L.brevis (+ vs -)"=~L_brevis,
                                     "A.thailandicus (+ vs -)"=~A_thai)) %>% 
  contrast("pairwise") %>% 
  map_dfr(~{confint(.) %>% data.frame},.id = "Bacteria")  
  

surv_oral_DCV_hazard_plot=ggplot(surv_oral_DCV_emm_bybac) +
  aes(x=Bacteria,y=estimate,ymin=asymp.LCL,ymax=asymp.UCL)+
  geom_bar(stat="identity")+
  geom_errorbar(width=.5)+
  theme_HMI()+
  labs(title="Cox regression coefficients",y=NULL,x="Contrast")
  
surv_oral_DCV_hazard_plot
```

##### Loads

###### Data loading

```
levels_oral_DCV
```

```
##     Infection Virus Wolbachia                Bacteria Replicate  fold_change
## 1        Oral   DCV     Wolb-          A.thailandicus         A 4.095451e-03
## 2        Oral   DCV     Wolb-          A.thailandicus         A 1.937841e+02
## 3        Oral   DCV     Wolb-          A.thailandicus         A 2.876064e+02
## 4        Oral   DCV     Wolb-          A.thailandicus         A 2.247753e-02
## 5        Oral   DCV     Wolb-          A.thailandicus         A 2.506705e-02
## 6        Oral   DCV     Wolb-          A.thailandicus         A 6.545664e-02
## 7        Oral   DCV     Wolb-          A.thailandicus         A 1.690198e-02
## 8        Oral   DCV     Wolb-          A.thailandicus         A 1.302040e+01
## 9        Oral   DCV     Wolb-          A.thailandicus         A 3.880886e-02
## 10       Oral   DCV     Wolb-          A.thailandicus         A 1.200334e+02
## 11       Oral   DCV     Wolb-          A.thailandicus         A 2.557181e-02
## 12       Oral   DCV     Wolb-          A.thailandicus         A 1.273942e+02
## 13       Oral   DCV     Wolb-          A.thailandicus         A 4.062609e-02
## 14       Oral   DCV     Wolb-          A.thailandicus         A 1.683767e-02
## 15       Oral   DCV     Wolb-          A.thailandicus         A 9.239409e-03
## 16       Oral   DCV     Wolb-          A.thailandicus         A 6.509972e+02
## 17       Oral   DCV     Wolb-          A.thailandicus         A 3.190251e+00
## 18       Oral   DCV     Wolb-          A.thailandicus         A 2.349923e+01
## 19       Oral   DCV     Wolb- A.thailandicus+L.brevis         A 4.414796e-01
## 20       Oral   DCV     Wolb- A.thailandicus+L.brevis         A 1.057841e-01
## 21       Oral   DCV     Wolb- A.thailandicus+L.brevis         A 2.607599e+02
## 22       Oral   DCV     Wolb- A.thailandicus+L.brevis         A 2.461421e+02
## 23       Oral   DCV     Wolb- A.thailandicus+L.brevis         A 1.575876e+01
## 24       Oral   DCV     Wolb- A.thailandicus+L.brevis         A 1.653396e-02
## 25       Oral   DCV     Wolb- A.thailandicus+L.brevis         A 6.047086e-02
## 26       Oral   DCV     Wolb- A.thailandicus+L.brevis         A 2.087750e+02
## 27       Oral   DCV     Wolb- A.thailandicus+L.brevis         A 3.255507e-02
## 28       Oral   DCV     Wolb- A.thailandicus+L.brevis         A 1.476763e-01
## 29       Oral   DCV     Wolb- A.thailandicus+L.brevis         A 3.404523e-01
## 30       Oral   DCV     Wolb- A.thailandicus+L.brevis         A 4.424875e+02
## 31       Oral   DCV     Wolb- A.thailandicus+L.brevis         A 9.451550e+01
## 32       Oral   DCV     Wolb- A.thailandicus+L.brevis         A 7.750015e-01
## 33       Oral   DCV     Wolb- A.thailandicus+L.brevis         A 4.102432e+02
## 34       Oral   DCV     Wolb- A.thailandicus+L.brevis         A 2.349508e+02
## 35       Oral   DCV     Wolb- A.thailandicus+L.brevis         A 2.384379e+02
## 36       Oral   DCV     Wolb- A.thailandicus+L.brevis         A 1.265507e-01
## 37       Oral   DCV     Wolb- A.thailandicus+L.brevis         A 2.673293e+02
## 38       Oral   DCV     Wolb- A.thailandicus+L.brevis         A 8.872622e+01
## 39       Oral   DCV     Wolb-                      GF         A 2.101305e+02
## 40       Oral   DCV     Wolb-                      GF         A 2.538929e-01
## 41       Oral   DCV     Wolb-                      GF         A 8.069758e+01
## 42       Oral   DCV     Wolb-                      GF         A 8.680283e-01
## 43       Oral   DCV     Wolb-                      GF         A 9.300758e+01
## 44       Oral   DCV     Wolb-                      GF         A 6.221297e+01
## 45       Oral   DCV     Wolb-                      GF         A 9.761453e+00
## 46       Oral   DCV     Wolb-                      GF         A 2.245334e+02
## 47       Oral   DCV     Wolb-                      GF         A 2.336538e+02
## 48       Oral   DCV     Wolb-                      GF         A 1.860078e+02
## 49       Oral   DCV     Wolb-                      GF         A 1.083842e-01
## 50       Oral   DCV     Wolb-                      GF         A 6.152647e-02
## 51       Oral   DCV     Wolb-                      GF         A 4.173224e+01
## 52       Oral   DCV     Wolb-                      GF         A 1.664506e-02
## 53       Oral   DCV     Wolb-                      GF         A 8.474724e+00
## 54       Oral   DCV     Wolb-                      GF         A 3.495036e-02
## 55       Oral   DCV     Wolb-                      GF         A 2.688568e-02
## 56       Oral   DCV     Wolb-                      GF         A 2.275210e+02
## 57       Oral   DCV     Wolb-                      GF         A 1.400366e-02
## 58       Oral   DCV     Wolb-                      GF         A 8.023574e-02
## 59       Oral   DCV     Wolb-                L.brevis         A 3.384012e-02
## 60       Oral   DCV     Wolb-                L.brevis         A 3.114575e-02
## 61       Oral   DCV     Wolb-                L.brevis         A 1.033118e+02
## 62       Oral   DCV     Wolb-                L.brevis         A 7.550458e+01
## 63       Oral   DCV     Wolb-                L.brevis         A 8.856482e-02
## 64       Oral   DCV     Wolb-                L.brevis         A 2.173975e+02
## 65       Oral   DCV     Wolb-                L.brevis         A 4.732113e+02
## 66       Oral   DCV     Wolb-                L.brevis         A 7.061812e-02
## 67       Oral   DCV     Wolb-                L.brevis         A 3.692491e+02
## 68       Oral   DCV     Wolb-                L.brevis         A 1.822449e+02
## 69       Oral   DCV     Wolb-                L.brevis         A 3.288235e+01
## 70       Oral   DCV     Wolb-                L.brevis         A 9.265187e+00
## 71       Oral   DCV     Wolb-                L.brevis         A 9.049559e-02
## 72       Oral   DCV     Wolb-                L.brevis         A 2.832156e+02
## 73       Oral   DCV     Wolb-                L.brevis         A 2.526813e+02
## 74       Oral   DCV     Wolb-                L.brevis         A 5.704622e+02
## 75       Oral   DCV     Wolb-                L.brevis         A 2.282273e+02
## 76       Oral   DCV     Wolb-                L.brevis         A 3.368895e+02
## 77       Oral   DCV     Wolb-                L.brevis         A 8.780943e+01
## 78       Oral   DCV     Wolb-                L.brevis         A 6.766212e-02
## 79       Oral   DCV     Wolb-          A.thailandicus         B 6.486837e+02
## 80       Oral   DCV     Wolb-          A.thailandicus         B 1.043416e+03
## 81       Oral   DCV     Wolb-          A.thailandicus         B 1.800421e-01
## 82       Oral   DCV     Wolb-          A.thailandicus         B 2.491226e+02
## 83       Oral   DCV     Wolb-          A.thailandicus         B 3.414707e-01
## 84       Oral   DCV     Wolb-          A.thailandicus         B 6.764768e+02
## 85       Oral   DCV     Wolb-          A.thailandicus         B 4.183205e-01
## 86       Oral   DCV     Wolb-          A.thailandicus         B 8.558934e+01
## 87       Oral   DCV     Wolb-          A.thailandicus         B 5.431571e+02
## 88       Oral   DCV     Wolb-          A.thailandicus         B 7.352695e+02
## 89       Oral   DCV     Wolb-          A.thailandicus         B 9.616050e+02
## 90       Oral   DCV     Wolb-          A.thailandicus         B 3.255010e+02
## 91       Oral   DCV     Wolb-          A.thailandicus         B 2.793241e-01
## 92       Oral   DCV     Wolb-          A.thailandicus         B 5.339670e+02
## 93       Oral   DCV     Wolb-          A.thailandicus         B 1.298040e+03
## 94       Oral   DCV     Wolb-          A.thailandicus         B 3.067618e+02
## 95       Oral   DCV     Wolb-          A.thailandicus         B 1.223441e+03
## 96       Oral   DCV     Wolb-          A.thailandicus         B 1.975268e-01
## 97       Oral   DCV     Wolb-          A.thailandicus         B 1.263679e+00
## 98       Oral   DCV     Wolb-          A.thailandicus         B 9.868570e+02
## 99       Oral   DCV     Wolb- A.thailandicus+L.brevis         B 6.350960e-02
## 100      Oral   DCV     Wolb- A.thailandicus+L.brevis         B 2.159932e-01
## 101      Oral   DCV     Wolb- A.thailandicus+L.brevis         B 1.907892e-01
## 102      Oral   DCV     Wolb- A.thailandicus+L.brevis         B 4.363025e+02
## 103      Oral   DCV     Wolb- A.thailandicus+L.brevis         B 3.930739e+02
## 104      Oral   DCV     Wolb- A.thailandicus+L.brevis         B 2.197452e-01
## 105      Oral   DCV     Wolb- A.thailandicus+L.brevis         B 4.617045e+01
## 106      Oral   DCV     Wolb- A.thailandicus+L.brevis         B 1.477395e+02
## 107      Oral   DCV     Wolb- A.thailandicus+L.brevis         B 7.706668e+02
## 108      Oral   DCV     Wolb- A.thailandicus+L.brevis         B 7.645305e+02
## 109      Oral   DCV     Wolb- A.thailandicus+L.brevis         B 1.435256e-01
## 110      Oral   DCV     Wolb- A.thailandicus+L.brevis         B 4.249582e+02
## 111      Oral   DCV     Wolb- A.thailandicus+L.brevis         B 3.241979e-01
## 112      Oral   DCV     Wolb- A.thailandicus+L.brevis         B 8.530657e+00
## 113      Oral   DCV     Wolb- A.thailandicus+L.brevis         B 2.111561e-01
## 114      Oral   DCV     Wolb- A.thailandicus+L.brevis         B 1.556189e+01
## 115      Oral   DCV     Wolb- A.thailandicus+L.brevis         B 7.503757e+02
## 116      Oral   DCV     Wolb- A.thailandicus+L.brevis         B 8.136802e+02
## 117      Oral   DCV     Wolb- A.thailandicus+L.brevis         B 5.673106e-01
## 118      Oral   DCV     Wolb- A.thailandicus+L.brevis         B 2.597690e+00
## 119      Oral   DCV     Wolb-                      GF         B 8.682047e+02
## 120      Oral   DCV     Wolb-                      GF         B 4.044086e-01
## 121      Oral   DCV     Wolb-                      GF         B 2.881676e-01
## 122      Oral   DCV     Wolb-                      GF         B 1.305669e-01
## 123      Oral   DCV     Wolb-                      GF         B 8.627023e+02
## 124      Oral   DCV     Wolb-                      GF         B 7.182405e-01
## 125      Oral   DCV     Wolb-                      GF         B 3.854878e-01
## 126      Oral   DCV     Wolb-                      GF         B 2.669800e-01
## 127      Oral   DCV     Wolb-                      GF         B 8.644690e+02
## 128      Oral   DCV     Wolb-                      GF         B 6.612567e-01
## 129      Oral   DCV     Wolb-                      GF         B 1.084761e+03
## 130      Oral   DCV     Wolb-                      GF         B 3.916688e+02
## 131      Oral   DCV     Wolb-                      GF         B 7.941073e+02
## 132      Oral   DCV     Wolb-                      GF         B 6.803100e+02
## 133      Oral   DCV     Wolb-                      GF         B 7.178272e-01
## 134      Oral   DCV     Wolb-                      GF         B 1.002716e+00
## 135      Oral   DCV     Wolb-                      GF         B 1.029595e+00
## 136      Oral   DCV     Wolb-                      GF         B 3.467792e-01
## 137      Oral   DCV     Wolb-                L.brevis         B 1.101976e+01
## 138      Oral   DCV     Wolb-                L.brevis         B 2.997392e-01
## 139      Oral   DCV     Wolb-                L.brevis         B 1.806750e-01
## 140      Oral   DCV     Wolb-                L.brevis         B 3.713659e+02
## 141      Oral   DCV     Wolb-                L.brevis         B 1.196609e+03
## 142      Oral   DCV     Wolb-                L.brevis         B 8.756857e+02
## 143      Oral   DCV     Wolb-                L.brevis         B 3.934015e+02
## 144      Oral   DCV     Wolb-                L.brevis         B 6.321630e+02
## 145      Oral   DCV     Wolb-                L.brevis         B 1.236049e-01
## 146      Oral   DCV     Wolb-                L.brevis         B 1.631198e-01
## 147      Oral   DCV     Wolb-                L.brevis         B 9.804772e-02
## 148      Oral   DCV     Wolb-                L.brevis         B 1.130397e-01
## 149      Oral   DCV     Wolb-                L.brevis         B 7.088015e+02
## 150      Oral   DCV     Wolb-                L.brevis         B 1.454319e-01
## 151      Oral   DCV     Wolb-                L.brevis         B 1.088681e+03
## 152      Oral   DCV     Wolb-                L.brevis         B 8.411722e-01
## 153      Oral   DCV     Wolb-                L.brevis         B 1.733656e-01
## 154      Oral   DCV     Wolb-                L.brevis         B 2.877985e+02
## 155      Oral   DCV     Wolb-                L.brevis         B 8.176309e+02
##     normalized_fold_change
## 1             4.491570e-04
## 2             2.125271e+01
## 3             3.154240e+01
## 4             2.465158e-03
## 5             2.749157e-03
## 6             7.178767e-03
## 7             1.853676e-03
## 8             1.427975e+00
## 9             4.256250e-03
## 10            1.316432e+01
## 11            2.804514e-03
## 12            1.397159e+01
## 13            4.455549e-03
## 14            1.846623e-03
## 15            1.013305e-03
## 16            7.139624e+01
## 17            3.498815e-01
## 18            2.577210e+00
## 19            4.841800e-02
## 20            1.160156e-02
## 21            2.859809e+01
## 22            2.699492e+01
## 23            1.728297e+00
## 24            1.813315e-03
## 25            6.631967e-03
## 26            2.289679e+01
## 27            3.570383e-03
## 28            1.619598e-02
## 29            3.733812e-02
## 30            4.852854e+01
## 31            1.036571e+01
## 32            8.499605e-02
## 33            4.499224e+01
## 34            2.576755e+01
## 35            2.614998e+01
## 36            1.387908e-02
## 37            2.931857e+01
## 38            9.730791e+00
## 39            2.304545e+01
## 40            2.784497e-02
## 41            8.850273e+00
## 42            9.519849e-02
## 43            1.020034e+01
## 44            6.823027e+00
## 45            1.070559e+00
## 46            2.462505e+01
## 47            2.562531e+01
## 48            2.039987e+01
## 49            1.188673e-02
## 50            6.747738e-03
## 51            4.576863e+00
## 52            1.825499e-03
## 53            9.294408e-01
## 54            3.833080e-03
## 55            2.948609e-03
## 56            2.495271e+01
## 57            1.535811e-03
## 58            8.799622e-03
## 59            3.711317e-03
## 60            3.415820e-03
## 61            1.133042e+01
## 62            8.280747e+00
## 63            9.713091e-03
## 64            2.384244e+01
## 65            5.189808e+01
## 66            7.744838e-03
## 67            4.049633e+01
## 68            1.998719e+01
## 69            3.606276e+00
## 70            1.016133e+00
## 71            9.924842e-03
## 72            3.106085e+01
## 73            2.771209e+01
## 74            6.256379e+01
## 75            2.503017e+01
## 76            3.694739e+01
## 77            9.630245e+00
## 78            7.420647e-03
## 79            7.538640e+02
## 80            1.212599e+03
## 81            2.092348e-01
## 82            2.895164e+02
## 83            3.968382e-01
## 84            7.861636e+02
## 85            4.861487e-01
## 86            9.946716e+01
## 87            6.312269e+02
## 88            8.544892e+02
## 89            1.117524e+03
## 90            3.782791e+02
## 91            3.246149e-01
## 92            6.205466e+02
## 93            1.508509e+03
## 94            3.565014e+02
## 95            1.421815e+03
## 96            2.295546e-01
## 97            1.468577e+00
## 98            1.146870e+03
## 99            7.380731e-02
## 100           2.510153e-01
## 101           2.217245e-01
## 102           5.070465e+02
## 103           4.568086e+02
## 104           2.553756e-01
## 105           5.365672e+01
## 106           1.716946e+02
## 107           8.956259e+02
## 108           8.884947e+02
## 109           1.667974e-01
## 110           4.938627e+02
## 111           3.767648e-01
## 112           9.913853e+00
## 113           2.453939e-01
## 114           1.808516e+01
## 115           8.720448e+02
## 116           9.456137e+02
## 117           6.592967e-01
## 118           3.018890e+00
## 119           1.008979e+03
## 120           4.699811e-01
## 121           3.348923e-01
## 122           1.517376e-01
## 123           1.002584e+03
## 124           8.346991e-01
## 125           4.479924e-01
## 126           3.102692e-01
## 127           1.004638e+03
## 128           7.684757e-01
## 129           1.260649e+03
## 130           4.551756e+02
## 131           9.228672e+02
## 132           7.906183e+02
## 133           8.342187e-01
## 134           1.165301e+00
## 135           1.196538e+00
## 136           4.030074e-01
## 137           1.280655e+01
## 138           3.483402e-01
## 139           2.099704e-01
## 140           4.315807e+02
## 141           1.390632e+03
## 142           1.017673e+03
## 143           4.571893e+02
## 144           7.346646e+02
## 145           1.436467e-01
## 146           1.895687e-01
## 147           1.139456e-01
## 148           1.313684e-01
## 149           8.237296e+02
## 150           1.690128e-01
## 151           1.265204e+03
## 152           9.775633e-01
## 153           2.014758e-01
## 154           3.344633e+02
## 155           9.502050e+02
```

###### Log Scale Model

```
levels_oral_DCV_model=lmer(log10(normalized_fold_change)~Bacteria+(1|Replicate),
                               data=levels_oral_DCV)
```

- Very skewed distribution of the residuals

```
resid(levels_oral_DCV_model) %>% qqnorm
```

- No effect of the gut microbiota

```
Anova(levels_oral_DCV_model)
```

```
## Analysis of Deviance Table (Type II Wald chisquare tests)
## 
## Response: log10(normalized_fold_change)
##           Chisq Df Pr(>Chisq)
## Bacteria 1.0288  3     0.7943
```

```
emmeans(levels_oral_DCV_model,pairwise~Bacteria)
```

```
## Note: Use 'contrast(regrid(object), ...)' to obtain contrasts of back-transformed estimates
```

```
## $emmeans
##  Bacteria                emmean    SE   df lower.CL upper.CL
##  GF                       0.270 0.838 1.19    -7.12     7.66
##  A.thailandicus           0.425 0.838 1.19    -6.96     7.81
##  L.brevis                 0.661 0.837 1.18    -6.80     8.12
##  A.thailandicus+L.brevis  0.474 0.836 1.17    -7.05     8.00
## 
## Degrees-of-freedom method: kenward-roger 
## Results are given on the log10 (not the response) scale. 
## Confidence level used: 0.95 
## 
## $contrasts
##  contrast                                   estimate    SE  df t.ratio p.value
##  GF - A.thailandicus                         -0.1547 0.392 150 -0.395  0.9790 
##  GF - L.brevis                               -0.3906 0.389 150 -1.004  0.7472 
##  GF - (A.thailandicus+L.brevis)              -0.2036 0.387 150 -0.527  0.9525 
##  A.thailandicus - L.brevis                   -0.2359 0.389 150 -0.606  0.9300 
##  A.thailandicus - (A.thailandicus+L.brevis)  -0.0488 0.387 150 -0.126  0.9993 
##  L.brevis - (A.thailandicus+L.brevis)         0.1870 0.384 150  0.487  0.9619 
## 
## Note: contrasts are still on the log10 scale 
## Degrees-of-freedom method: kenward-roger 
## P value adjustment: tukey method for comparing a family of 4 estimates
```

```
emmeans(levels_oral_DCV_model,~Bacteria) %>% 
  cld(Letters=letters)
```

```
## Note: Use 'contrast(regrid(object), ...)' to obtain contrasts of back-transformed estimates
```

```
##  Bacteria                emmean    SE   df lower.CL upper.CL .group
##  GF                       0.270 0.838 1.19    -7.12     7.66  a    
##  A.thailandicus           0.425 0.838 1.19    -6.96     7.81  a    
##  A.thailandicus+L.brevis  0.474 0.836 1.17    -7.05     8.00  a    
##  L.brevis                 0.661 0.837 1.18    -6.80     8.12  a    
## 
## Degrees-of-freedom method: kenward-roger 
## Results are given on the log10 (not the response) scale. 
## Confidence level used: 0.95 
## Note: contrasts are still on the log10 scale 
## P value adjustment: tukey method for comparing a family of 4 estimates 
## significance level used: alpha = 0.05
```

###### Kruskal-Wallis

- No effect of the gut microbiota

```
levels_oral_DCV_kw=kruskal.test(normalized_fold_change~Bacteria,data=levels_oral_DCV)

levels_oral_DCV_kw
```

```
## 
##  Kruskal-Wallis rank sum test
## 
## data:  normalized_fold_change by Bacteria
## Kruskal-Wallis chi-squared = 0.93079, df = 3, p-value = 0.818
```

```
kruskalmc(normalized_fold_change~Bacteria,data=levels_oral_DCV)
```

```
## Multiple comparison test after Kruskal-Wallis 
## p.value: 0.05 
## Comparisons
##                                         obs.dif critical.dif difference
## GF-A.thailandicus                      7.421053     27.16926      FALSE
## GF-L.brevis                            9.373144     26.99454      FALSE
## GF-A.thailandicus+L.brevis             5.327632     26.82750      FALSE
## A.thailandicus-L.brevis                1.952092     26.99454      FALSE
## A.thailandicus-A.thailandicus+L.brevis 2.093421     26.82750      FALSE
## L.brevis-A.thailandicus+L.brevis       4.045513     26.65053      FALSE
```

###### Plot

```
levels_oral_DCV_plot=ggplot(levels_oral_DCV)+
  aes(x=Bacteria,y=normalized_fold_change)+
  geom_beeswarm(cex = 2,aes(shape=Bacteria,color=Bacteria))+
   scale_color_manual(values=c("darkgray","black","black","black"))+
  scale_y_log10(labels=prettyNum)+
  guides(shape="none",color="none")+
  facet_grid(~"")+
  theme_HMI()+
  stat_summary(fun="median",geom="crossbar",width=.5)+
  labs(title="Relative DCV levels",y=NULL)

levels_oral_DCV_plot
```

##### Arranged plot

```
fig3_plotlist=list(surv_oral_DCV_panel,
  surv_oral_DCV_hazard_plot+facet_grid(~""),
  levels_oral_DCV_plot%+%filter(levels_oral_DCV,Replicate=="A") )

 
fig3=wrap_plots(fig3_plotlist,guides="collect")+
  patchwork::plot_annotation(tag_levels = "A")&
  theme(legend.position = "bottom",plot.tag = element_text(face="bold"))

fig3
```

```
ggsave2(filename = here("manuscript","fig3.png"),plot = fig3,
        width=18,
        height=6,
        units="cm",scale=1.5)
```

#### Session info

```
sessioninfo::session_info()
```

```
## - Session info ---------------------------------------------------------------
##  setting  value                       
##  version  R version 4.0.3 (2020-10-10)
##  os       Windows 10 x64              
##  system   x86_64, mingw32             
##  ui       RTerm                       
##  language (EN)                        
##  collate  English_Europe.1252         
##  ctype    English_Europe.1252         
##  tz       Europe/London               
##  date     2021-02-18                  
## 
## - Packages -------------------------------------------------------------------
##  package      * version date       lib source        
##  abind          1.4-5   2016-07-21 [1] CRAN (R 4.0.0)
##  assertthat     0.2.1   2019-03-21 [1] CRAN (R 4.0.0)
##  backports      1.2.0   2020-11-02 [1] CRAN (R 4.0.3)
##  bdsmatrix    * 1.3-4   2020-01-13 [1] CRAN (R 4.0.0)
##  beeswarm       0.2.3   2016-04-25 [1] CRAN (R 4.0.0)
##  boot           1.3-25  2020-04-26 [1] CRAN (R 4.0.0)
##  broom        * 0.7.3   2020-12-16 [1] CRAN (R 4.0.3)
##  car          * 3.0-10  2020-09-29 [1] CRAN (R 4.0.3)
##  carData      * 3.0-4   2020-05-22 [1] CRAN (R 4.0.0)
##  cellranger     1.1.0   2016-07-27 [1] CRAN (R 4.0.0)
##  class          7.3-17  2020-04-26 [1] CRAN (R 4.0.2)
##  classInt       0.4-3   2020-04-07 [1] CRAN (R 4.0.2)
##  cli            2.2.0   2020-11-20 [1] CRAN (R 4.0.3)
##  coda           0.19-4  2020-09-30 [1] CRAN (R 4.0.3)
##  codetools      0.2-18  2020-11-04 [1] CRAN (R 4.0.3)
##  colorspace     2.0-0   2020-11-11 [1] CRAN (R 4.0.3)
##  cowplot      * 1.1.1   2020-12-30 [1] CRAN (R 4.0.3)
##  coxme        * 2.2-16  2020-01-14 [1] CRAN (R 4.0.0)
##  crayon         1.3.4   2017-09-16 [1] CRAN (R 4.0.0)
##  curl           4.3     2019-12-02 [1] CRAN (R 4.0.0)
##  data.table     1.13.6  2020-12-30 [1] CRAN (R 4.0.3)
##  DBI            1.1.1   2021-01-15 [1] CRAN (R 4.0.3)
##  dbplyr         2.0.0   2020-11-03 [1] CRAN (R 4.0.2)
##  deldir         0.2-9   2021-01-16 [1] CRAN (R 4.0.3)
##  digest         0.6.27  2020-10-24 [1] CRAN (R 4.0.3)
##  dplyr        * 1.0.3   2021-01-15 [1] CRAN (R 4.0.3)
##  e1071          1.7-4   2020-10-14 [1] CRAN (R 4.0.3)
##  ellipsis       0.3.1   2020-05-15 [1] CRAN (R 4.0.2)
##  emmeans      * 1.5.3   2020-12-09 [1] CRAN (R 4.0.3)
##  estimability   1.3     2018-02-11 [1] CRAN (R 4.0.0)
##  evaluate       0.14    2019-05-28 [1] CRAN (R 4.0.0)
##  expm           0.999-6 2021-01-13 [1] CRAN (R 4.0.3)
##  fansi          0.4.2   2021-01-15 [1] CRAN (R 4.0.3)
##  farver         2.0.3   2020-01-16 [1] CRAN (R 4.0.0)
##  forcats      * 0.5.0   2020-03-01 [1] CRAN (R 4.0.0)
##  foreign        0.8-81  2020-12-22 [1] CRAN (R 4.0.3)
##  fs             1.5.0   2020-07-31 [1] CRAN (R 4.0.2)
##  gdata          2.18.0  2017-06-06 [1] CRAN (R 4.0.0)
##  generics       0.1.0   2020-10-31 [1] CRAN (R 4.0.3)
##  ggbeeswarm   * 0.6.0   2017-08-07 [1] CRAN (R 4.0.0)
##  ggplot2      * 3.3.3   2020-12-30 [1] CRAN (R 4.0.3)
##  ggpubr       * 0.4.0   2020-06-27 [1] CRAN (R 4.0.2)
##  ggsignif       0.6.0   2019-08-08 [1] CRAN (R 4.0.0)
##  glue           1.4.2   2020-08-27 [1] CRAN (R 4.0.3)
##  gmodels        2.18.1  2018-06-25 [1] CRAN (R 4.0.2)
##  gridExtra      2.3     2017-09-09 [1] CRAN (R 4.0.0)
##  gtable         0.3.0   2019-03-25 [1] CRAN (R 4.0.0)
##  gtools         3.8.2   2020-03-31 [1] CRAN (R 4.0.0)
##  haven          2.3.1   2020-06-01 [1] CRAN (R 4.0.2)
##  here         * 1.0.1   2020-12-13 [1] CRAN (R 4.0.3)
##  highr          0.8     2019-03-20 [1] CRAN (R 4.0.0)
##  hms            1.0.0   2021-01-13 [1] CRAN (R 4.0.3)
##  htmltools      0.5.0   2020-06-16 [1] CRAN (R 4.0.2)
##  httr           1.4.2   2020-07-20 [1] CRAN (R 4.0.2)
##  jsonlite       1.7.1   2020-09-07 [1] CRAN (R 4.0.3)
##  KernSmooth     2.23-18 2020-10-29 [1] CRAN (R 4.0.3)
##  km.ci          0.5-2   2009-08-30 [1] CRAN (R 4.0.0)
##  KMsurv         0.1-5   2012-12-03 [1] CRAN (R 4.0.0)
##  knitr          1.30    2020-09-22 [1] CRAN (R 4.0.3)
##  labeling       0.4.2   2020-10-20 [1] CRAN (R 4.0.3)
##  lattice        0.20-41 2020-04-02 [2] CRAN (R 4.0.3)
##  LearnBayes     2.15.1  2018-03-18 [1] CRAN (R 4.0.0)
##  lifecycle      0.2.0   2020-03-06 [1] CRAN (R 4.0.0)
##  lme4         * 1.1-26  2020-12-01 [1] CRAN (R 4.0.3)
##  lubridate      1.7.9.2 2020-11-13 [1] CRAN (R 4.0.3)
##  magrittr     * 2.0.1   2020-11-17 [1] CRAN (R 4.0.3)
##  maptools       1.0-2   2020-08-24 [1] CRAN (R 4.0.3)
##  MASS         * 7.3-53  2020-09-09 [1] CRAN (R 4.0.3)
##  Matrix       * 1.2-18  2019-11-27 [1] CRAN (R 4.0.3)
##  minqa          1.2.4   2014-10-09 [1] CRAN (R 4.0.0)
##  modelr         0.1.8   2020-05-19 [1] CRAN (R 4.0.2)
##  multcomp     * 1.4-15  2020-11-14 [1] CRAN (R 4.0.3)
##  multcompView   0.1-8   2019-12-19 [1] CRAN (R 4.0.0)
##  munsell        0.5.0   2018-06-12 [1] CRAN (R 4.0.0)
##  mvtnorm      * 1.1-1   2020-06-09 [1] CRAN (R 4.0.0)
##  nlme           3.1-151 2020-12-10 [1] CRAN (R 4.0.3)
##  nloptr         1.2.2.2 2020-07-02 [1] CRAN (R 4.0.2)
##  openxlsx       4.2.3   2020-10-27 [1] CRAN (R 4.0.3)
##  patchwork    * 1.1.1   2020-12-17 [1] CRAN (R 4.0.3)
##  pbkrtest       0.5-0.1 2020-12-18 [1] CRAN (R 4.0.3)
##  pgirmess     * 1.6.9   2018-03-12 [1] CRAN (R 4.0.3)
##  pillar         1.4.7   2020-11-20 [1] CRAN (R 4.0.3)
##  pkgconfig      2.0.3   2019-09-22 [1] CRAN (R 4.0.0)
##  plyr           1.8.6   2020-03-03 [1] CRAN (R 4.0.0)
##  purrr        * 0.3.4   2020-04-17 [1] CRAN (R 4.0.0)
##  R6             2.5.0   2020-10-28 [1] CRAN (R 4.0.3)
##  raster         3.4-5   2020-11-14 [1] CRAN (R 4.0.3)
##  Rcpp           1.0.5   2020-07-06 [1] CRAN (R 4.0.2)
##  readr        * 1.4.0   2020-10-05 [1] CRAN (R 4.0.3)
##  readxl         1.3.1   2019-03-13 [1] CRAN (R 4.0.0)
##  reprex         0.3.0   2019-05-16 [1] CRAN (R 4.0.0)
##  rgdal          1.5-19  2021-01-05 [1] CRAN (R 4.0.3)
##  rgeos          0.5-5   2020-09-07 [1] CRAN (R 4.0.3)
##  rio            0.5.16  2018-11-26 [1] CRAN (R 4.0.0)
##  rlang          0.4.10  2020-12-30 [1] CRAN (R 4.0.3)
##  rmarkdown      2.6     2020-12-14 [1] CRAN (R 4.0.3)
##  rprojroot      2.0.2   2020-11-15 [1] CRAN (R 4.0.3)
##  rstatix        0.6.0   2020-06-18 [1] CRAN (R 4.0.2)
##  rstudioapi     0.13    2020-11-12 [1] CRAN (R 4.0.3)
##  rvest          0.3.6   2020-07-25 [1] CRAN (R 4.0.2)
##  sandwich       3.0-0   2020-10-02 [1] CRAN (R 4.0.3)
##  scales         1.1.1   2020-05-11 [1] CRAN (R 4.0.2)
##  sessioninfo    1.1.1   2018-11-05 [1] CRAN (R 4.0.0)
##  sf             0.9-7   2021-01-06 [1] CRAN (R 4.0.3)
##  sp             1.4-5   2021-01-10 [1] CRAN (R 4.0.3)
##  spData         0.3.8   2020-07-03 [1] CRAN (R 4.0.2)
##  spdep          1.1-5   2020-06-29 [1] CRAN (R 4.0.2)
##  splancs        2.01-40 2017-04-16 [1] CRAN (R 4.0.3)
##  statmod        1.4.35  2020-10-19 [1] CRAN (R 4.0.3)
##  stringi        1.4.6   2020-02-17 [1] CRAN (R 4.0.0)
##  stringr      * 1.4.0   2019-02-10 [1] CRAN (R 4.0.0)
##  survival     * 3.2-7   2020-09-28 [1] CRAN (R 4.0.3)
##  survminer    * 0.4.8   2020-07-25 [1] CRAN (R 4.0.2)
##  survMisc       0.5.5   2018-07-05 [1] CRAN (R 4.0.0)
##  TH.data      * 1.0-10  2019-01-21 [1] CRAN (R 4.0.0)
##  tibble       * 3.0.5   2021-01-15 [1] CRAN (R 4.0.3)
##  tidyr        * 1.1.2   2020-08-27 [1] CRAN (R 4.0.3)
##  tidyselect     1.1.0   2020-05-11 [1] CRAN (R 4.0.2)
##  tidyverse    * 1.3.0   2019-11-21 [1] CRAN (R 4.0.2)
##  units          0.6-7   2020-06-13 [1] CRAN (R 4.0.2)
##  utf8           1.1.4   2018-05-24 [1] CRAN (R 4.0.0)
##  vctrs          0.3.6   2020-12-17 [1] CRAN (R 4.0.3)
##  vipor          0.4.5   2017-03-22 [1] CRAN (R 4.0.0)
##  withr          2.4.0   2021-01-16 [1] CRAN (R 4.0.3)
##  xfun           0.19    2020-10-30 [1] CRAN (R 4.0.3)
##  xml2           1.3.2   2020-04-23 [1] CRAN (R 4.0.2)
##  xtable         1.8-4   2019-04-21 [1] CRAN (R 4.0.0)
##  yaml           2.2.1   2020-02-01 [1] CRAN (R 4.0.0)
##  zip            2.1.1   2020-08-27 [1] CRAN (R 4.0.3)
##  zoo            1.8-8   2020-05-02 [1] CRAN (R 4.0.2)
## 
## [1] C:/Users/Nelson/Documents/R/win-library/4.0
## [2] C:/Program Files/R/R-4.0.3/library
```

##### PDF for figures

```
ggsave2(here("manuscript","fig1.pdf"),
                 plot = fig1,
                 width = 18,height = 9,units = "cm",scale=1.5)

ggsave2(here("manuscript","fig2.pdf"),
        plot = fig2,width=14,height=6,units="cm",scale=1.5) 

ggsave2(here("manuscript","fig3.pdf"),
        plot = fig3,
        width=18,height=6,units="cm",scale=1.5) 

ggsave2(here("manuscript","figS1.pdf"),
        plot=fig_s1,
        width = 14,height=9,units="cm",scale=1.5)
```
